# A Murine Model of Lyme Disease Demonstrates That *Borrelia burgdorferi* Colonizes the Dura Mater and Induces Inflammation in the Central Nervous System

**DOI:** 10.1101/2020.04.24.060681

**Authors:** Timothy Casselli, Ali Divan, Yvonne Tourand, Heidi L. Pecoraro, Catherine A. Brissette

## Abstract

Lyme disease, which is caused by infection with *Borrelia burgdorferi* and related species, can lead to inflammatory pathologies affecting the joints, heart, and nervous systems including the central nervous system (CNS). Inbred laboratory mice are effective models for characterizing *B. burgdorferi* infection kinetics and host immune responses in joints and heart tissues; however, similar studies are lacking in the CNS of these animals. Here we characterize the kinetics of *B. burgdorferi* colonization and associated immune responses in the CNS of infected C3H mice during early and subacute infection. *B. burgdorferi* colonized the dura mater following needle or tick challenge, and induced expression of inflammatory cytokines and a robust IFN response as well as histopathological changes. A sterile IFN response in the absence of *B. burgdorferi* or inflammatory cytokines was unique to the brain parenchyma, and could provide insights into the mechanism of inflammatory CNS pathology associated with this important pathogen.

## INTRODUCTION

Lyme disease (LD), which is caused by infection with the bacterial pathogen *Borrelia burgdorferi* and related species, is a prevalent and continually emerging vector-borne disease throughout North America and Europe (Hinckley et al., 2014; Schwartz, Hinckley, Mead, Hook, & Kugeler, 2017). Disseminated infection can lead to a number of subacute and persistent inflammatory pathologies affecting the joints (arthritis), heart (carditis and heart block), and nervous systems including the central nervous system (CNS) (Bratton, Whiteside, Hovan, Engle, & Edwards, 2008; Halperin, 2015; Wormser et al., 2006). CNS manifestations can include lymphocytic meningitis, radiculoneuritis, and cranial neuritis. More serious complications during late persistent infection can include encephalitis, and vasculitis (Halperin, 2015).

Inbred laboratory mice have served as effective models to characterize *B. burgdorferi* infection kinetics, as well as the host immune responses involved in pathogen burden control and inflammatory pathology. After initial challenge by either needle inoculation or tick transmission, spirochetes disseminate throughout the animal and colonize peripheral tissues including skin, joints, and heart (S. Barthold, Persing, Armstrong, & Peeples, 1991; Zeidner, Schneider, Dolan, & Piesman, 2001). Peripheral tissue colonization is persistent in the absence of antibiotic treatment. Infection of disease-susceptible C3H mice results in subacute arthritis and carditis characterized by joint swelling and histopathological manifestations similar to human disease including leukocyte infiltration (Armstrong, Barthold, Persing, & Beck, 1992; S. W. Barthold, Beck, Hansen, Terwilliger, & Moody, 1990).

Previous studies using a variety of laboratory mouse backgrounds and co-culture models have provided a detailed picture of the differential roles of the host immune responses to infection. *B. burgdorferi* and its components can induce host cell production of innate and T cell-mediated inflammatory cytokines as well as chemokines for monocytes, polymorphonuclear leukocytes (PMNs), and lymphocytes both in vitro and during murine infection (Casselli et al., 2017; Crandall et al., 2006; Schramm et al., 2012; Verhaegh, Joosten, & Oosting, 2017). The innate inflammatory cytokine response is mediated largely through Toll-like receptor (TLR)-2/MyD88/TRIF signaling and NF-κB (Bolz et al., 2004; Ebnet, Brown, Siebenlist, Simon, & Shaw, 1997; Hirschfeld et al., 1999; Petnicki-Ocwieja et al., 2013; Wooten et al., 2002; Wooten, Modur, McIntyre, & Weis, 1996), which is essential for efficient control of spirochete burden in conjunction with a lymphocyte-mediated adaptive anti-*B. burgdorferi* response (S. W. Barthold, Sidman, & Smith, 1992; Bolz et al., 2004; Brown & Reiner, 1999b; Wooten et al., 2002). In contrast, inflammatory pathology does not require TLR-2 signaling, B cells, or T cells, but is perpetuated by a robust early interferon (IFN) response in disease-susceptible mice (S. W. Barthold et al., 1992; Bolz et al., 2004; Brown & Reiner, 1999b; Crandall et al., 2006; Miller et al., 2008; Wooten et al., 2002). Interestingly, both type I and type II interferons contribute to the early induction of IFN-stimulated genes (ISGs); however type II IFN and STAT1 are dispensable for murine arthritis development, whereas blockage of type I IFN leads to reduced joint pathology (Brown, Blaho, Fritsche, & Loiacono, 2006; Brown & Reiner, 1999a; Miller et al., 2008). The role of IFN signaling in murine Lyme carditis is less well understood; however, the signals required for disease manifestation appear to be distinct in the heart and joint (Brown et al., 2006; Lochhead et al., 2014; Olson et al., 2009).

Despite the utility of the murine model for investigating Lyme arthritis and carditis, similar studies on the kinetics of *B. burgdorferi* colonization and the resulting host responses are lacking in the CNS of these animals. Given the neurological sequelae associated with *B. burgdorferi* infection in addition to arthritis and carditis in Lyme disease patients, a tractable animal model for understanding host-pathogen interactions in the CNS is needed (Garcia-Monco & Benach, 2013). Previously, we reported that *B. burgdorferi* colonizes the dura mater of C3H mice during late disseminated infection, with an associated increase in dura T cell numbers (Divan et al., 2018). In the current study, we characterized *B. burgdorferi* colonization kinetics in the dura mater during early and subacute infection (7 and 28 days post-infection, respectively), as well as the associated host immune responses in the dura mater and brain parenchyma, as these timepoints have been demonstrated to most consistently replicate inflammatory pathology in murine models of Lyme disease (Armstrong et al., 1992; S. W. Barthold et al., 1990).

Overall, we report that *B. burgdorferi* routinely colonizes the meninges in laboratory mice during early and subacute infection, and induces similar localized inflammatory gene expression profiles as other peripheral tissues as well as histopathological changes. A sterile IFN response in the absence of *B. burgdorferi* or inflammatory cytokines is unique to the brain parenchyma, and could provide insights into the mechanism of inflammatory CNS pathology associated with this important pathogen.

## RESULTS

### *B. burgdorferi* colonize the dura mater during early and subacute infection

Based on our previous finding that *B. burgdorferi* strain 297 (Bb_297) colonized the dura mater of C3H mice during late disseminated infection (75 days post-infection) (Divan et al., 2018), we set out to determine the kinetics of dura mater colonization during early and subacute infection. Mice were initially infected by intradermal inoculation with 10^6^ bacteria for 3-28 days, after which the dura were isolated and assayed by fluorescent immunohistochemistry (f-IHC) to identify the presence, location, and quantity of spirochetes. Dura stained with anti-*B. burgdorferi* antibodies from uninfected mice or 3-day infected mice did not show the presence of intact spirochetes, whereas spirochetes were identified at all later infection timepoints (Figure 1). Spirochetes were ubiquitous throughout the dura mater (Figure 1A), and were largely not associated with nearby blood vessels. Bacterial burden peaked at day 7 post-infection, and declined by days 14 and 28 to levels similar to that seen at day 75 post-infection (Figure 1B) (Divan et al., 2018). Multiphoton imaging of the dura mater of mice infected with Green Fluorescent Protein-expressing Bb_297 (Bb_297-GFP) confirmed spirochetes are extravascular, alive, and motile (Movie 1). Despite the abundance of Bb_297-GFP in the dura mater, no spirochetes were observed in the brain parenchyma (not shown), indicating that spirochetes were limited to the meninges in the CNS of mice.

**Figure 1.**
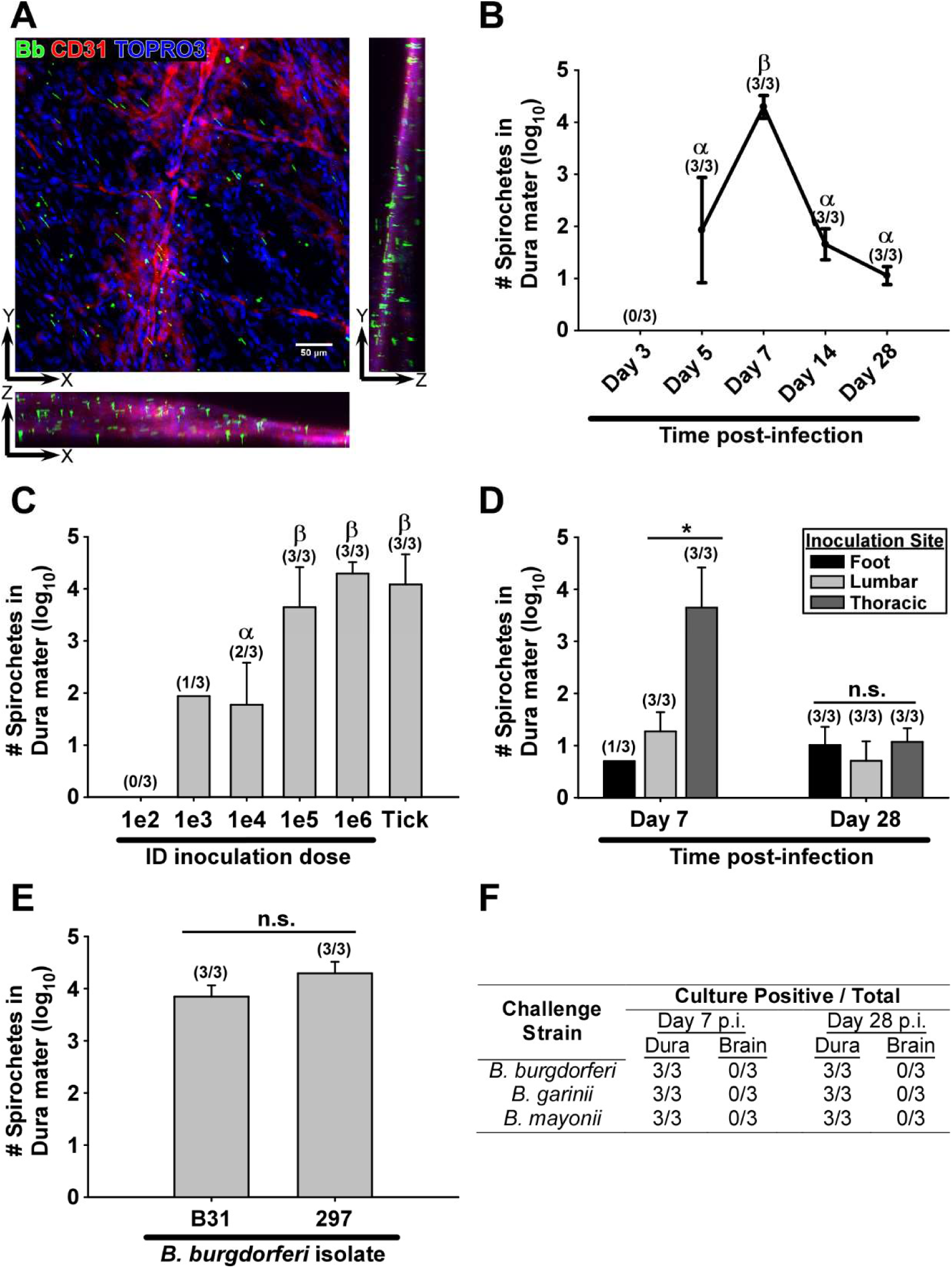
*B. burgdorferi* infects the dura ubiquitously during early infection. **A.** Representative confocal z-series showing *B. burgdorferi* (Bb, green), blood vessels (CD31, red), and nucleated cells (TOPRO3, blue) in the dura mater of C3H mice on day 7 of infection with 10^6^ spirochetes. X-axis rotation (XZ) and Y-axis rotation (ZY) of image are also shown. **B.** Number of intact spirochetes in the dura mater (log_10_ mean ± s.d.) at various time points after infection with 10^6^ spirochetes. Symbols α and β indicate statistically different groups (p ≤ 0.003; α = 0.05) as determined by one‐way ANOVA followed by all pairwise multiple comparison (Holm‐Sidak). **C.** Number of spirochetes in dura mater (log_10_ mean ± s.d.) on day 7 of infection using increasing needle inoculation doses as well as tick transmission. Symbols α and β indicate statistically different groups (0.021 < p < 0.048; α = 0.05) as determined by one‐way ANOVA followed by all pairwise multiple comparison (Holm‐Sidak). **D.** *B. burgdorferi* burden in the dura (log_10_ mean ± s.d.) based on site of inoculation (foot, lumbar, thoracic), and duration of infection (day 7, day 28) of mice infected intradermally with 10^6^ spirochetes. Asterisk indicates significant difference as determined using Student’s t-test (p = 0.009; α = 0.05). “n.s” denotes no statistical significance by one-way ANOVA (p = 0.4; α = 0.05). **E.** Bacterial burden in the dura on day 7 of infection after inoculation with 10^6^ *B. burgdorferi* strain B31, or strain 297. “n.s” denotes no statistical significance by Student’s t-test (p = 0.07; α = 0.05). In **(B-E)**, Spirochetes were counted in isolated dura mater by systematic counting using f-IHC, epifluorescence microscopy, and gridded coverslips, as described in methods. Number of dura samples with detectable spirochetes / total n are listed for each group above the bar. Conditions with less than two positive dura samples were omitted from statistical analysis. **F.** Number of isolated dura samples resulting in positive cultures in BSK medium from mice infected for 7-28 days with *B. burgdorferi* strain 297, *B. garinii*, or *B. mayonii* as determined using darkfield microscopy.

Challenge dose is an important consideration during experimental infection of laboratory animals. As our initial experiments were done using 10^6^ bacteria, we repeated the experiment with decreasing challenge doses of Bb_297. At day 7 post-infection, no spirochetes were observed in the dura mater of mice challenged with 10^2^ bacteria, whereas challenge doses of 10^3^ and 10^4^ spirochetes resulted in only 1/3 and 2/3 positive dura, respectively, with relatively low levels of detectable spirochetes (Figure 1C). All mice had detectable spirochetes in the dura mater at day 7 post-infection after challenge doses from 10^5^-10^6^ bacteria, with higher bacterial burdens compared to the lower challenge doses (Figure 1C).

Given the dose-dependent nature of dura colonization by needle inoculation, we repeated the experiment using mice infected by nymphal tick transmission to determine the relevance after exposure from the natural vector. Seven days post-transmission feeding, dura spirochete burdens in all mice were similar to that seen using a needle inoculation dose of 10^6^ bacteria (Figure 1C). Therefore, all future experiments were carried out using intradermal needle challenge doses of 10^6^ spirochetes to mimic dura colonization observed after tick transmission.

We also examined the effects of inoculation site on dura colonization. Mice were infected by needle inoculation in either the footpad, dorsal lumbar skin, or dorsal thoracic skin. At day 7 post-infection there was a marked difference in dura colonization efficiency between inoculation sites, with higher spirochete burdens observed after infection at sites more proximal to the dura mater (Figure 1D). This difference was abrogated by day 28 post-infection, indicating that only the initial peak in burden is dependent on inoculation site, and not the ability to persistently colonize this tissue.

Bb_297 is a clinical isolate from the CSF of a patient diagnosed with neuroborreliosis (Steere et al., 1983). To determine if additional Lyme disease *Borrelia* isolates are capable of dura colonization in mice, we repeated the experiment with the *B. burgdorferi* tick isolate B31 (Burgdorfer et al., 1982). No difference was seen in bacterial burden in the dura mater between *B. burgdorferi* sensu stricto (s.s.) isolates (Figure 1E). We were also able to culture both *B. garinii* (a European *B. burgdorferi* sensu lato (s.l.) species) as well as *B. mayonii* (a North American *B. burgdorferi* s.l. species) from the dura mater of perfused mice at both day 7 and day 28 post-infection; however, brain cultures from all animals were negative (Kriuchechnikov, Korenberg, Shcherbakov, Kovalevskii Iu, & Levin, 1988; Pritt et al., 2016). Taken together, colonization of the dura mater in mice is a general phenomenon for clinical and tick isolates of *B. burgdorferi* s.s., as well as both North American and European isolates of *B. burgdorferi* s.l.

### *B. burgdorferi* infection leads to leukocyte infiltration in the dura mater

A hallmark of laboratory murine Lyme arthritis and carditis is a subacute leukocytic infiltration between 2-4 weeks post-infection (Armstrong et al., 1992; S. W. Barthold et al., 1990). We examined the dura mater and brain parenchyma at 7 days post-infection (during peak spirochetemia in the meninges) as well as 28 days post-infection (timepoint of peak Lyme arthritis) for signs of inflammatory cell infiltrate.

Dura sections isolated by craniotomy and stained with hematoxylin/eosin revealed perivascular leukocyte infiltrate and/or mild/minimal meningitis without vascular hemorrhage consisting of mainly mononuclear cells with rare PMNs in 5/6 infected mice, with the exception of one 7-day infected mouse (Figure 2A). None of the control mice showed signs of dura leukocyte infiltration. Additionally, more than half of all dura sections from 7-day infected mice showed signs of perivascular infiltrate associated with vascular hemorrhage (34.0±2.6 sections; 57%), compared to 5.0±2.6 sections (8%) and 5.6±5.0 (9%) of uninfected and 28-day infected dura, respectively, indicating a decrease in vascular integrity at day 7 post-infection (Figure 2A).

**Figure 2.**
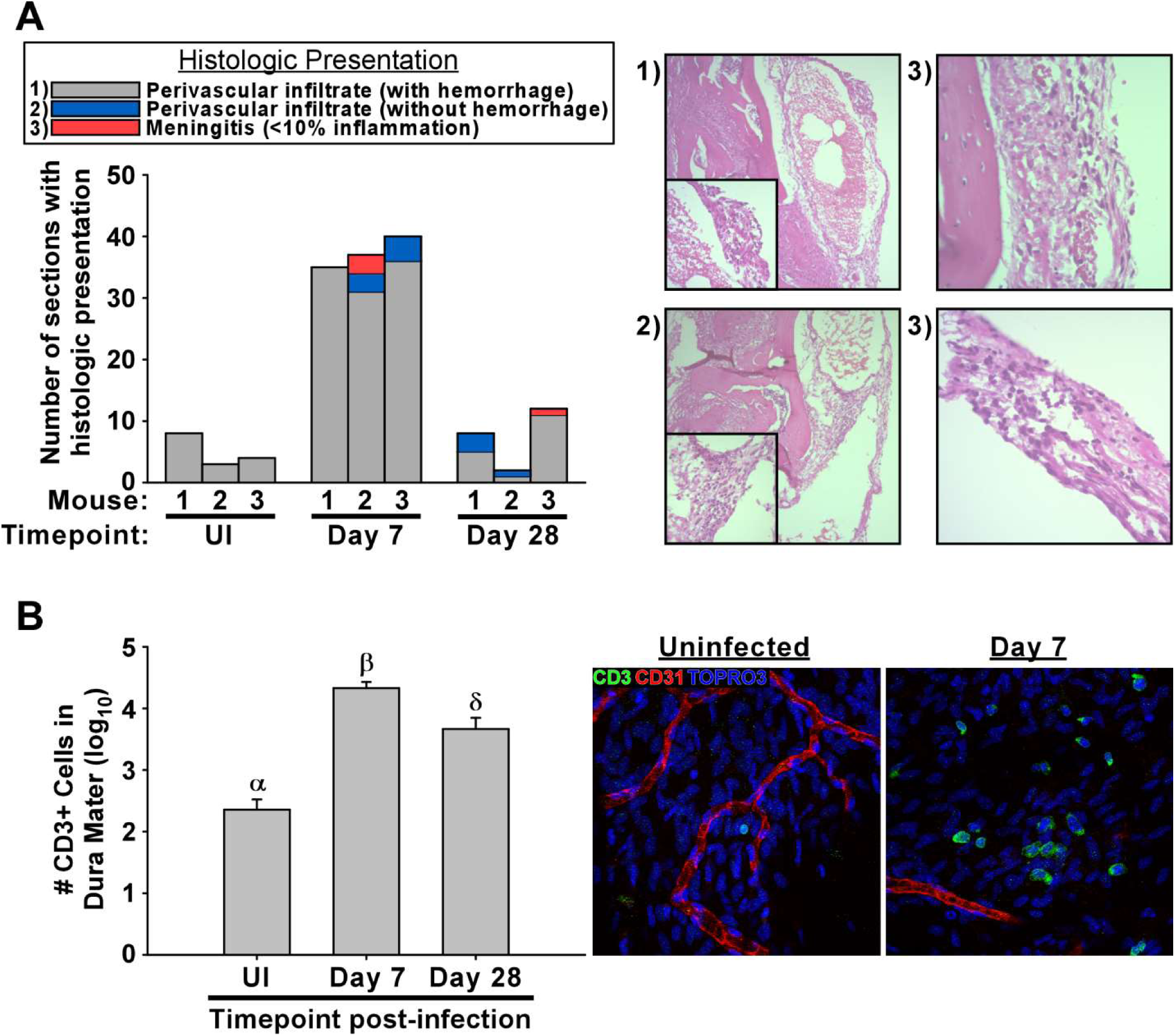
Infection with *B. burgdorferi* leads to leukocyte infiltration including T cells. **A.** Skullcaps with attached dura were fixed, and 60 representative coronal sections were systematically evaluated for inflammation by histopathology. Stacked bargraph shows number of sections from each mouse with histologic presentations as shown in the legend. Timepoint of infection is shown for each mouse. Representative images from sections with severity scores of 1 (perivascular infiltrate associated with hemorrhage; 100X magnification, inset 400X magnification), 2 (perivascular infiltrate without hemorrhage; 100X magnification, inset 400X magnification), and 3 (minimal meningitis; 400X magnification) are shown. **B.** Bar graph showing number of CD3+ cells ± s.d. in the dura mater from day 7 and 28 post-infection compared to uninfected controls (n=3). Symbols α, β, and δ indicate statistically different groups (p < 0.006; α = 0.05) as determined by one‐way ANOVA followed by all pairwise multiple comparison (Holm‐Sidak). Representative confocal images are shown from uninfected and 7-day infected mice. T cells (CD3, green), blood vessels (CD31, red), and nucleated cells (TOPRO3, blue) are shown.

Similar histopathologic analysis of brain sections did not reveal leukocytic infiltration of the cerebral cortex, hippocampus, and thalamus in either the control or infected mice; consistent with the lack of detectable spirochetes in the parenchyma. Meningeal congestion was occasionally observed in all mice, including control mice, but perivascular hemorrhage admixed with leukocytes was observed only in the infected mice. In brain sections, perivascular hemorrhage was characterized by extravasated red blood cells admixed with rare to few mononuclear cells expanding the meninges, especially along the ventral cortex, and choroid plexus or extending into the subependyma of the lateral ventricle (Figure 3). Interestingly, leukocyte infiltrates associated with hemorrhage were seen in nearly all (99.4%) of the 176 brain sections evaluated on day 7 infected mice compared to 74% (129/174) of the day 28 infected mice.

**Figure 3.**
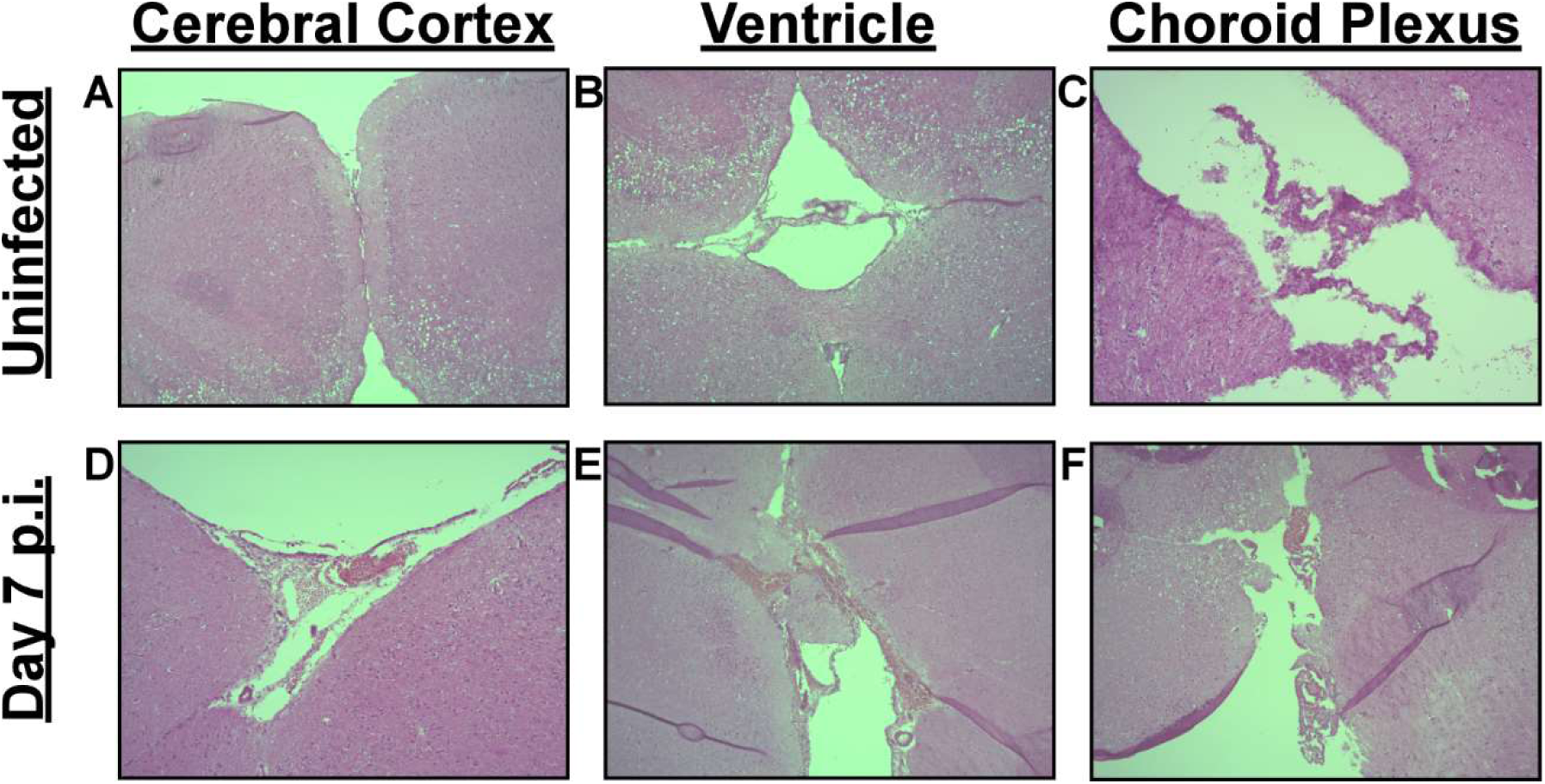
Infection with *B. burgdorferi* is associated with perivascular hemorrhage admixed with leukocytes, without leukocytic infiltration into the brain parenchyma. Uninfected mice showed no microscopic evidence of inflammation or hemorrhage in brain sections examined **(A-C)**, while perivascular hemorrhage with attendant leukocytes was observed in the cerebral meninges **(D)**, ventricles extending into the subependymal zone **(E)**, and throughout the choroid plexus **(F)**. Leukocytic infiltration was not noted in the cerebral cortex, hippocampus, or thalamus in either control or infected mice.

We previously showed dura colonization by *B. burgdorferi* during late disseminated infection is associated with an increase in T lymphocytes (Divan et al., 2018). We therefore assessed the number of CD3+ cells in the dura mater at day 7 and day 28 post-infection by f-IHC. Examination of uninfected dura showed the presence of scattered CD3+ cells that appeared small and spherical (Figure 2B). At days 7 and 28 post-infection, there was a marked increase in the number of CD3+ cells, which appeared larger and had morphology/staining profiles consistent with more active/motile cells.

Taken together, dura colonization by *B. burgdorferi* is associated with a localized infiltration of leukocytes including T cells. Additionally, increases in perivascular hemorrhage associated with blood vessels of the dura mater, cerebral meninges, and ventricles, suggest a breakdown of vascular integrity; however, this does not appear to be sufficient to allow detectable infiltration of leukocytes or *B. burgdorferi* into the brain parenchyma.

### *B. burgdorferi* infection is associated with large scale changes in gene expression in the dura mater and brain parenchyma

#### Overview of gene expression analysis

Studies on gene expression changes in joint and heart tissues have provided insights into the localized host responses to *B. burgdorferi* colonization (Crandall et al., 2006; Kelleher Doyle et al., 1998); however, gene expression changes in the CNS of mice during *B. burgdorferi* infection remain unclear. Therefore, we took an unbiased approach to examine potential changes in the dura mater as well as the brain cortex and hippocampus at day 7 post-infection using RNA sequencing (RNA-seq). Prior to euthanasia, mice were anesthetized and perfused to remove gene expression signatures from circulating blood.

Over 1,900 and 1,400 genes were significantly upregulated or downregulated in the dura mater of infected mice, respectively (Fold-Change ≥ 1.5, padj ≤ 0.05, basemean > 20) (Figure 4; see supplemental Table S1 for complete comparative expression list and individual padj values). Additionally, brain cortex and hippocampus both exhibited a substantial number of differentially expressed genes (DEGs; 258 and 158, respectively) (Figure 4; supplemental Tables S2-S3). Although the number of upregulated and downregulated DEGs in the dura mater were comparable (56% vs 44% upregulated vs downregulated, respectively), the majority of DEGs in the brain were upregulated (93% in cortex, 82% in hippocampus). These responses were consistent across biological treatments and replicates (supplemental Figure S1). Collectively, these data show a robust response to *B. burgdorferi* colonization in the dura mater as well as changes in gene expression in the brain parenchyma at day 7 post-infection, despite the absence of a local presence of spirochetes or infiltrating leukocytes in the brain.

**Figure 4.**
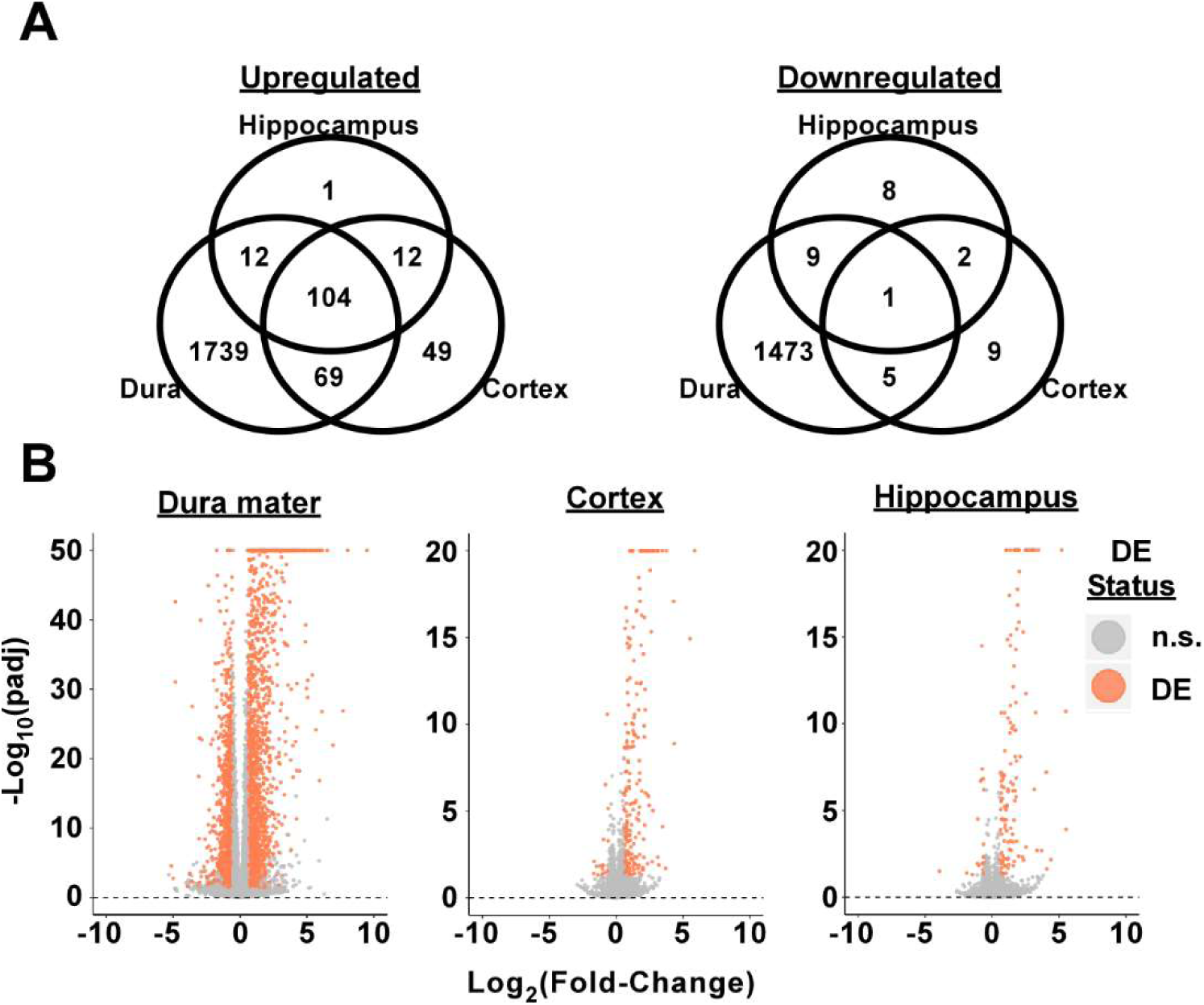
Infection with *B. burgdorferi* leads to transcriptome changes in the meninges and brain parenchyma. **A.** Venn diagram showing the number of distinct and common significantly upregulated and downregulated genes in the dura mater, brain cortex, and hippocampus (padj ≤ 0.05, basemean ≥ 20, fold-change ≥ 1.5) of 7-day infected mice compared to uninfected controls (n=4 per timepoint). **B.** Volcano plots from the three tissues tested comparing -log(padj) to log_2_(fold-change) for all genes. Red color indicates DEGs, while gray indicates genes not significantly different between infected and control mice (n.s.). See supplemental Figure S1 for principle component analysis and hierarchical clustering of individual samples.

#### U pregulated genes in the dura mater, cortex, and hippocampus demonstrate profiles consistent with IFN response, whereas upregulation of inflammatory cytokines and chemokines are restricted to the dura mater

The top five most enriched molecular function gene ontology (GO) terms from genes upregulated in the dura mater all pertained to cytokine/chemokine production, activity, or their receptors (supplemental Figure S2A). In contrast, GO terms enriched in upregulated genes in both the cortex and hippocampus were related to antigen processing and presentation, double-stranded RNA-binding, and GTPase activity (supplemental Figure S2B-C). When only those genes commonly upregulated in all three tissues were considered, a similar GO profile was observed as cortex or hippocampus alone, indicating that these molecular functions were enriched across all tissues tested (supplemental Figure S2D).

Examination of individual upregulated genes associated with the five most enriched GO terms in the dura mater revealed a number of hallmark inflammatory cytokines induced by *B. burgdorferi* including *tnf*, *il6*, and *il1β* (Figure 5A-C; Table S1). Several chemokines of monocytes (*ccl2*, *ccl7*), PMNs (*cxcl1*, *cxcl2*), and lymphocytes (*ccl2*, *ccl7*, *ccl19*, *cxcl9*, *cxcl10*, *cxcl13*) were also upregulated. The majority of these genes were upregulated greater than 10-fold in the dura mater, however were not differentially expressed in the RNA-seq dataset in the cortex or hippocampus (Figure 5C; Tables S1-S3). As an inflammatory cytokine response is thought to be initiated in response to *B. burgdorferi* antigens through TLR signaling and NF-κB activation (Bolz et al., 2004; Ebnet et al., 1997; Hirschfeld et al., 1999; Petnicki-Ocwieja et al., 2013; Wooten et al., 2002; Wooten et al., 1996), it is perhaps not surprising that upregulation of these genes is largely limited to the dura mater where spirochete burden is prominent. Indeed, we observed significant upregulation of TLR genes *tlr1*, *tlr2*, *tlr6*, *tlr7*, *tlr8*, and *tlr9* as well as adaptor molecules including *myd88* (Table S1), and Signaling Pathway Impact Analysis (SPIA) revealed activation of both TLR (pGFWER = 1.30E-06) and NF-κB (pGFWER = 1.15E-16) signalling pathways exclusively in the dura mater (supplemental Figures S3-S4).

**Figure 5.**
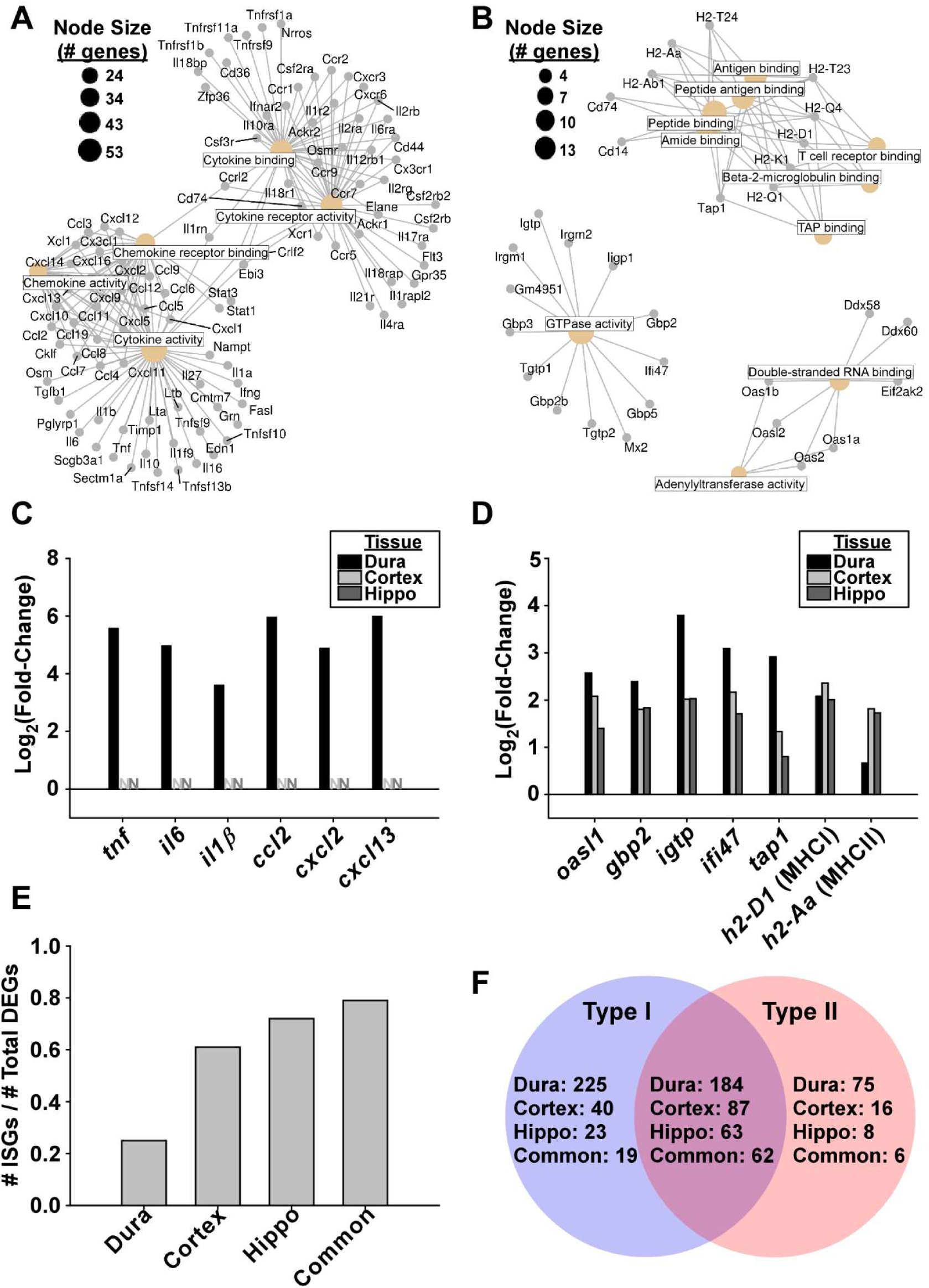
Upregulated genes in the dura mater, cortex, and hippocampus demonstrate profiles consistent with IFN response, whereas upregulation of inflammatory cytokines are restricted to the dura mater. **A.** Cnet plot (generated using ClusterProfiler (Yu et al., 2012)) showing individual genes associated with the top 5 GO terms upregulated in the RNA-seq dataset from the dura mater of infected C3H mice. Colored nodes represent GO terms, with node descriptions represented as boxed text. Node size correlates with the number of associated genes, as shown in the legend. Individual upregulated genes are represented by gray dots, with lines connecting to the relevant GO terms. **B.** Cnet plot as described in **(A)**, showing top 10 enriched GO terms and associated genes commonly upregulated in all three tissues. **C.** Barplot representation of log_2_(fold-change) from RNA-seq data of selected genes from **(A)**. Bars denote magnitude of change for DEGs, whereas “N” represents genes not differentially expressed, demonstrating the tissue specificity of cytokine response. **D.** Barplot representation from RNA-seq data of selected genes from **(B)**, showing similarity of response in the three tissues. **E.** Proportion of upregulated genes from each tissue as well as those genes commonly upregulated in all tissues that are predicted to be upregulated by IFN (Rusinova et al., 2013). **F.** Venn diagram showing number of upregulated DEGs from each tissue predicted to be stimulated by Type I or Type II interferons (Rusinova et al., 2013).

A subset of 104 genes were found to be similarly upregulated in all three tissues tested. Examination of these common upregulated genes associated with the 10 most enriched GO terms revealed a number of IFN-inducible GTPases and signaling molecules including *oasl1*/*2*, *gbp2*/*3*/*5*, *igtp*, as well as several genes involved in antigen processing/presentation by both MHCI and MHCII (e.g. *H2-Aa*, *H2-D1*, *H2-K1*, *H2-Q1*, *tap1*) (Figure 5B for gene expression; supplemental Figure S5 for SPIA). In fact, the majority of upregulated genes in the cortex (61%), hippocampus (72%), and those common to all three tissues (79%) were associated with IFN response, while ISGs constituted 25% of the total number of upregulated genes in the dura mater (Rusinova et al., 2013)(Figure 5E). ISGs associated with both type I and type II IFN were identified in all tested tissues (Figure 5F). Intriguingly, type II IFN was only detected at low levels in the dura mater, and type I IFN was not detectable in any tissue type despite a robust IFN response; although genes coding for both type I and type II interferon receptors were significantly upregulated in the dura mater (Table S1).

Gene expression profiles from the dura mater were consistent with our histopathological findings of an influx of monocytic immune cells including T cells. Expression of endothelial cell adhesion molecules *vcam1* and *icam1* were significantly upregulated in the dura of infected mice (Table S1). Additionally, SPIA showed activation of the T cell receptor signalling pathway (pGFWER = 8.48E-05) including increased expression of all three CD3 subunits as well as downstream molecules (supplemental Table S1; Figure S6), whereas no increase was observed for the B cell marker *cd19*. Expression of *itgam* (CD11b) was highly upregulated in infected dura (log_2_(fold-change) = 3.66; padj = 1e-50), as well as genes encoding Ly6C (*ly6c1*, *ly6c2*), while *itgax* (CD11c) and Ly6G genes were not significantly altered, consistent with an increase in monocytes/macrophages.

In addition to increased expression of MHC genes in the brain parenchyma, both *itgax* (CD11c) and Ly6c genes were differentially upregulated in the cortex of infected mice, while *itgam* (CD11b) expression levels were only modestly increased (45% upregulation; padj = 0.02). Hippocampus samples showed an upregulation of Ly6c genes in response to infection; however, no difference was detected in expression levels of CD11b or CD11c genes. No significant differences in B cell or T cell marker genes were detected in the brain parenchyma, consistent with a lack of observed leukocytes in the parenchyma as determined by histopathologic analysis.

Collectively these data show the presence of two distinct immune profiles in the CNS of infected mice. In the dura mater, the presence of *B. burgdorferi* is associated with upregulation of genes consistent with TLR/NF-κB signalling and associated inflammatory cytokines and chemokines in addition to a robust IFN response. In contrast, the brain parenchyma exhibits mainly an IFN response to *B. burgdorferi* infection without an associated cytokine response, despite a lack of detectable spirochetes in these tissues.

#### Colonization of the dura mater is associated with altered expression of genes associated with the extracellular matrix, cell-adhesion, and wound repair

Biological process GO term enrichment from downregulated genes in the dura mater of infected mice show genes involved in ECM organization, connective tissue development, cell-substrate adhesion, and wound repair (Figure 6A). Examination of individual genes showed significant downregulation of several ECM components including fibrillary collagens (*col1a1*, *col1a2*, *col2a1*, *col3a1*, *col5a1*, *col5a2*, *col5a3*, *col6a2*, *col6a3*, *col9a1*, *col9a2*, *col11a1*, *col11a2*, *col15a1 col16a1*), basement membrane (BM)-associated collagens (*col4a5*, *col4a6*, *col8a1*, *col8a2*), and transmembrane collagens (*col13a1*, *col24a1*, *col25a1*). Other BM components including perivascular BM-associated lamanins (*lama2*, *lama3*, *lamb1*, *lamb3*), nidogen (*nid1*, *nid2*), heparin sulfate proteoglycans (perlecan; *hspg2*) and other ECM components (fibrillin; *fbn1*, *fbn2*) were also downregulated in infected dura (Figure 6B; supplementary Table S1). Genes coding for the ECM proteins fibronectin (*fn1*) and decorin (*dcn*) were highly expressed in the dura mater; however, expression levels were not altered in response to infection (supplementary Table S1).

**Figure 6.**
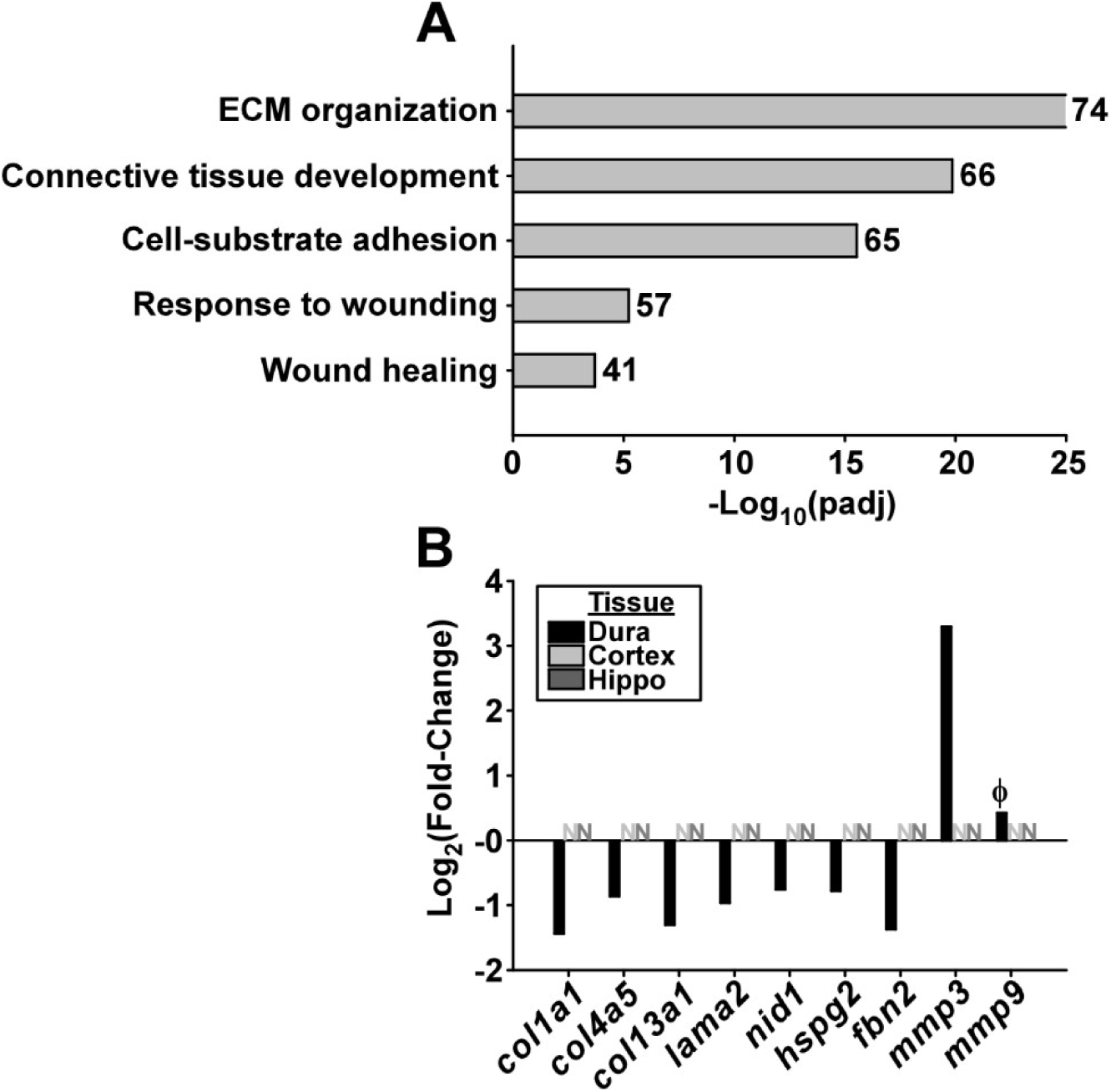
Colonization of the dura mater is associated with downregulation of genes associated with the extracellular matrix. **A.** Selected biological process GO terms enriched in downregulated genes in the dura mater. Numbers to the right of horizontal bars show the number of downregulated DEGs associated with each term. Bar size represents significance of enrichment (-log(padj)). **B.** Log_2_(fold-change) of selected genes from RNA-seq dataset associated with GO terms from **(A).** Bars represent magnitude of change for differentially expressed genes, while “N” represents genes not significantly differentially expressed in the RNA-seq dataset; demonstrating the tissue specificity of the response. Φ represents statistical significance (p = 0.003), but does not meet the pre-determined fold-change cut-off to be classified as a DEG (fold-change = 1.3; DEG cut-off = 1.5).

The above ECM components affected by *B. burgdorferi* infection are the targets of host proteases that allow for tissue remodelling under physiologic or pathologic conditions (Sbardella et al., 2012). The matrix metalloproteinase gene *mmp3* was significantly upregulated in the dura mater of infected mice (Figure 6B), which is involved in tissue reorganization, wound repair, and activation of other MMPs (e.g. MMP9) (Sbardella et al., 2012), and has previously shown to be induced by *B. burgdorferi* in C3H mice (Behera, Hildebrand, Scagliotti, Steere, & Hu, 2005; Crandall et al., 2006). The murine Lyme arthritis-associated *mmp9* was also significantly upregulated in the dura mater of infected mice at day 7 post-infection (Crandall et al., 2006; Heilpern et al., 2009; Hu et al., 2001); however, the magnitude of expression change did not reach our arbitrary fold-change cut-off for differential expression calling (padj = 0.003; Fold-change = 1.3; DEG cut-off = 1.5).

Taken together, these data illustrate large-scale downregulation of genes coding for structural proteins in the dura mater in response to *B. burgdorferi* infection, as well as induction of the proteases that target their degradation/reorganization.

#### Increased expression of inflammatory cytokines persists during disseminated infection, whereas IFN response is limited to early infection

Host gene expression profiles from the RNA-seq dataset at day 7 post-infection in the dura mater and brain parenchyma were confirmed by qRT-PCR of representative genes. Additionally, the presence of live *B. burgdorferi* was confirmed by qRT-PCR of the bacterial *flaB* gene. Additional timepoints from 7-56 days post-infection were assessed to determine the kinetics of host responses to infection. The *tnf* gene was chosen as a proxy for inflammatory cytokines, as well as *gbp2* as proxy for ISGs. C*xcl10* expression was also assayed, as this gene has been demonstrated as an ISG, with evidence of non-interferon induction by NF-κB (Brownell et al., 2014).

*FlaB* transcripts were readily detected at all timepoints in the dura mater, tibiotarsal joints, and heart tissues, whereas no *flaB* transcript was detectable in either the cortex or hippocampus at any timepoint (Figure 7), consistent with our initial microscopy and culture data. Likewise, *tnf* levels were elevated in the dura mater, joint, and heart tissues at all timepoints compared to uninfected controls, but were not increased in the cortex or hippocampus, supporting our RNA-seq data showing increased cytokine expression correlates with the presence of spirochetes. In contrast, the ISG *gbp2* was elevated in all tissues tested including brain tissues at day 7 post-infection, again in agreement with the RNA-seq dataset. Expression of *gbp2* returned to uninfected levels by day 28 post-infection in all tissues, which is a pattern that has been previously reported for ISGs in joints during murine Lyme arthritis (Crandall et al., 2006). Cxcl10 was found to be elevated in all tissues at day 7 post-infection, consistent with its induction by IFN. This gene remained elevated in dura, joint, and heart tissue up to 56 days post-infection, however returned to uninfected levels after day 7 post-infection in the brain. This profile of *cxcl10* expression suggests that in addition to early induction in all tissues as part of an IFN response, the persistence of spirochetes in peripheral tissues and the dura mater is sufficient for sustained elevated expression of this gene.

**Figure 7.**
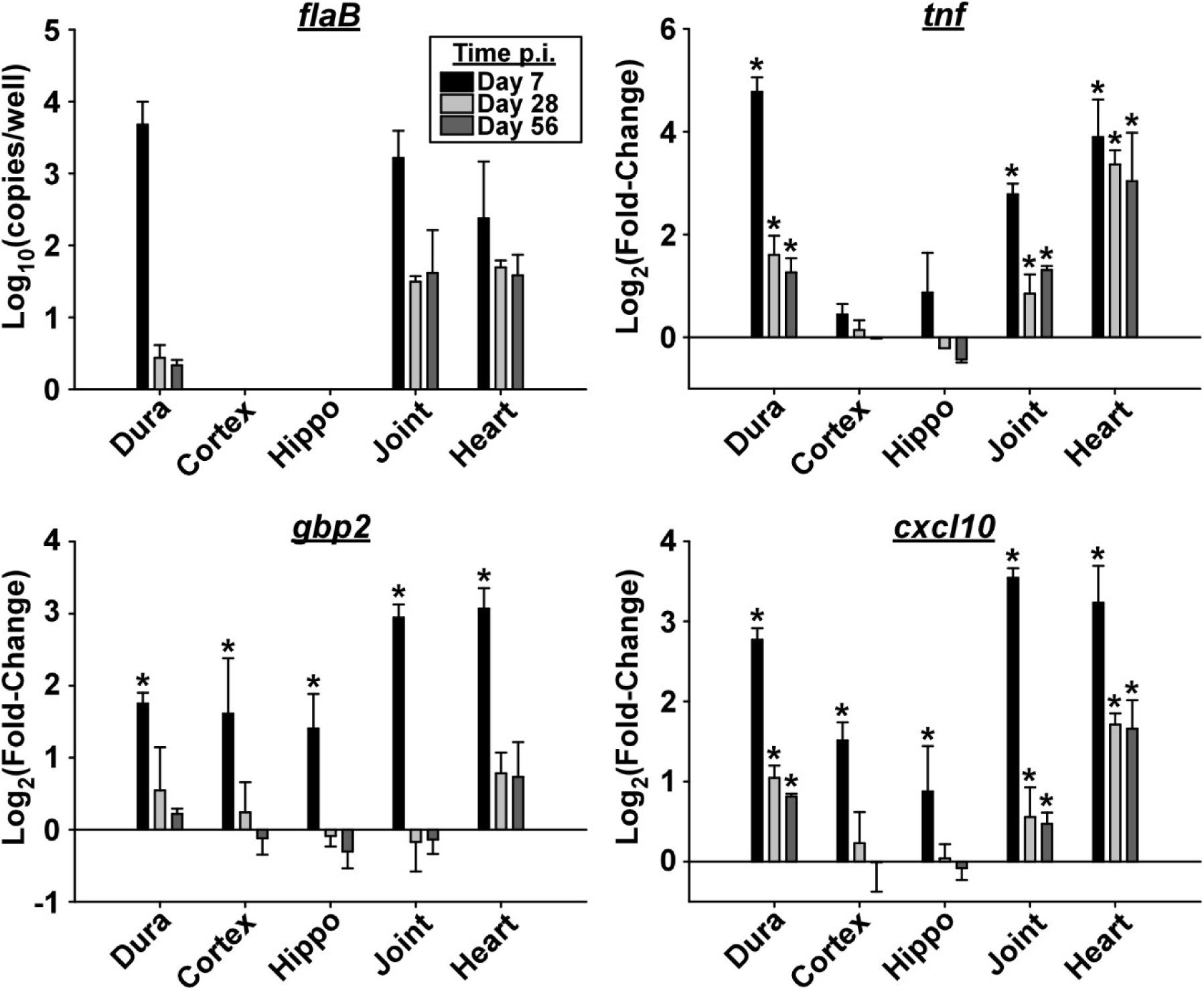
Increased expression of inflammatory cytokines persists during late disseminated infection, whereas IFN response is limited to early infection. qRT-PCR of selected genes from dura mater (dura), cortex, hippocampus (hippo), tibiotarsal joint (joint), and heart tissues from day 7-56 post-infection as shown in the legend (n=4 per timepoint). The *B. burgdorferi* gene *flaB* is displayed as gene copies per reaction (log_10_ mean ± s.d.) as determined using standard curves. Levels of *flaB* copies were below the limit of detection in the cortex and hippocampus. Host genes *tnf*, *gbp2*, and *cxcl10* are displayed as log_2_(fold-change) (mean ± s.d.) compared to uninfected controls as determined using the ∆∆Ct method. Asterisks indicate statistical significance (0.001 ≤ p ≤ 0.026; α = 0.05) using one-way ANOVA followed by Dunnett’s test.

These data show that *B. burgdorferi* persistently colonize the dura mater, and induces a sustained inflammatory cytokine response in addition to an early transient IFN response. In contrast, *B. burgdorferi* do not colonize the brain parenchyma; however, a transient IFN response is still induced in the cortex and hippocampus of infected animals that returned to uninfected levels as infection persisted.

### T cells and B cells are necessary for efficient reduction of *B. burgdorferi* load during persistent infection in the dura mater, however are not required for leukocyte infiltration or upregulation of ISGs

Studies using immunodeficient SCID and *rag*-/- mice have clearly demonstrated a role for B cells and T cells in controlling *B. burgdorferi* burden in peripheral tissues during persistent infection; however, these cells are not required for early induction of ISGs or the development of murine Lyme arthritis (S. W. Barthold et al., 1992; Bolz et al., 2004; Miller et al., 2008). Similarly, Bb_297 burden was comparable in the dura mater of wildtype C3H and SCID mice at day 7 post-infection, whereas bacterial numbers were elevated in SCID mice at days 14 and 28 post-infection (Figure 8A), indicating a role for adaptive immunity in dura spirochete control during persistent infection. Histopathologic analysis of the dura mater from infected C3H-SCID showed a similar pattern of leukocyte infiltration as observed in wildtype mice in response to *B. burgdorferi* infection, with more prevalent perivascular leukocyte infiltrate at day 7 post-infection compared to similarly treated C3H mice (Figure 8B; see also Figure 2A). Additionally, T cells and B cells were not required for induction of ISGs, as levels of both *gbp2* and *cxcl10* were elevated in infected SCID mice compared to uninfected controls in the dura, cortex, hippocampus, and tibiotarsal joints (Figure 8C-D), similar to levels seen in wild-type mice (Figure 7).

**Figure 8.**
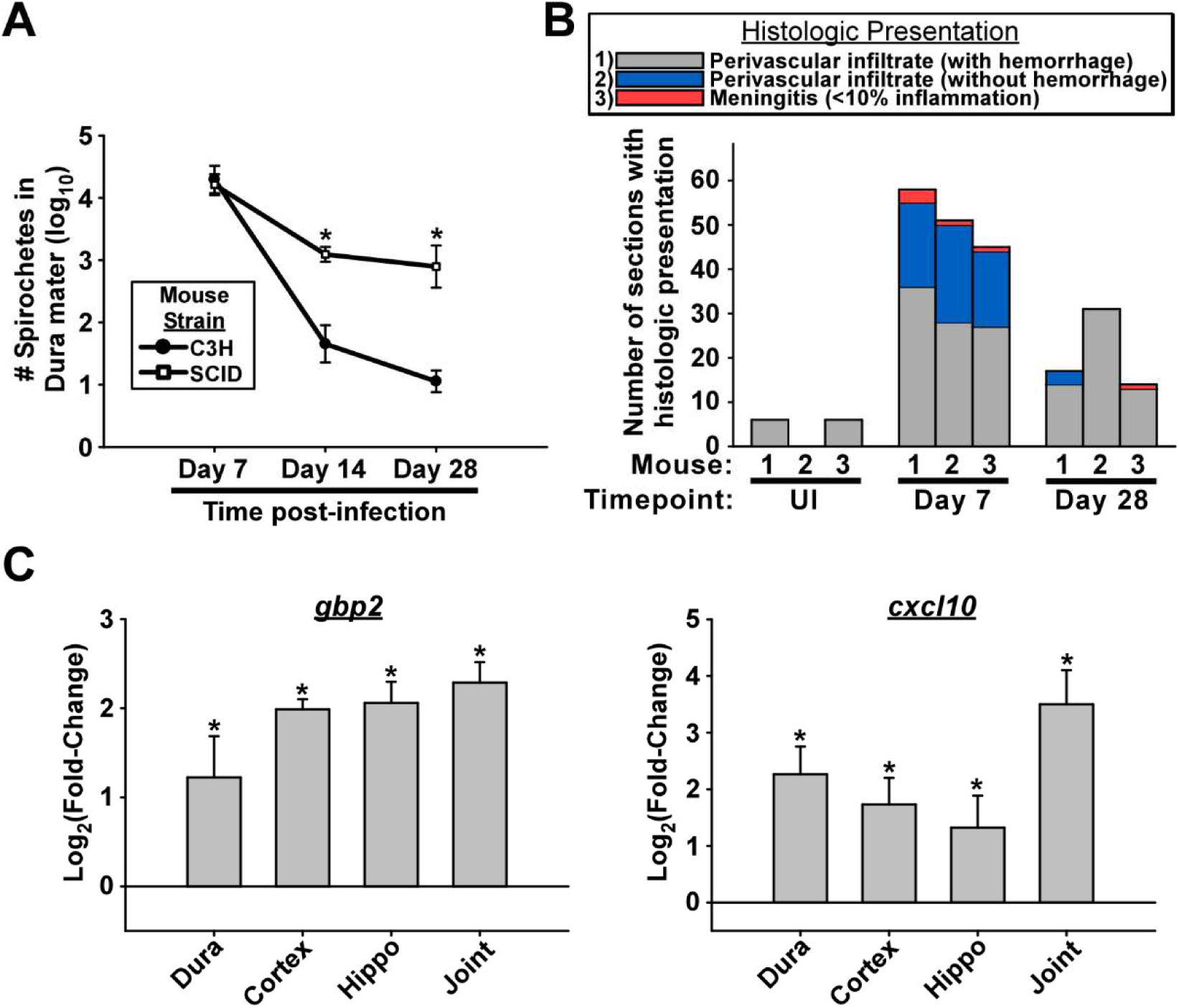
B cells and T cells are required for controlling spirochete burden during persistent infection, however are not required for leukocyte infiltration or upregulation of ISGs. **A.** Bacterial burden (log_10_ mean ± s.d.) in the dura mater of C3H vs C3H-SCID mice from day 7-28 post-infection (n=3 per condition). Spirochetes were counted in isolated dura mater by systematic counting using f-IHC, epifluorescence microscopy, and gridded coverslips, as described in methods. Asterisks indicate statistical significance (p ≤ 0.001; α = 0.05) using Student’s t-test. **B.** Histopathology of dura mater from C3H-SCID mice at days 7-28 post-infection and uninfected controls (see detailed description in the legend of Figure 2). Stacked bargraph shows number of sections (out of 60) from each mouse with histologic presentations as shown in the legend. Timepoint of infection is shown for each mouse. **C.** qRT-PCR of indicated ISGs from dura mater (dura), cortex, hippocampus (hippo), and tibiotarsal joint (joint) of C3H-SCID mice at day 7 post-infection (n=3). Bars are displayed as log_2_(fold-change) ± s.d. compared to uninfected control samples for each tissue as determined using the ∆∆Ct method. Asterisks indicate statistical significance (0.001 ≤ p ≤ 0.04; α = 0.05) using Student’s t-test.

Collectively, these data show a similar requirement for T cells and B cells for pathogen control and IFN response in the CNS as previously reported in peripheral tissues during infection with *B. burgdorferi* (S. W. Barthold et al., 1992; Bolz et al., 2004; Miller et al., 2008).

## DISCUSSION

### The dura mater is a site of *Borrelia* colonization in mice during early and subacute infection

We report here for the first time, the kinetics of *B. burgdorferi* colonization of the meninges as well as the CNS host response to infection in a tractable laboratory animal model of Lyme disease. Following infection of mice by either needle inoculation or tick transmission, *B. burgdorferi* readily colonized the extravascular dura mater, reaching peak burdens of over 10^4^ bacteria at 7 days post-infection. Although epidemiological evidence suggest that some Lyme disease *Borrelia* may be more neurotropic than others (Pachner & Steiner, 2007), we did not find evidence of this in our model of dura colonization. The similar distribution of spirochetes in the dura mater between isolates Bb_B31 and Bb_297 suggests that ribosomal spacer type (RST) and OspC type are not determinants of dura colonization among *B. burgdorferi* s.s. isolates (G. Wang, van Dam, Schwartz, & Dankert, 1999). We were also able to readily culture both North American and European *B. burgdorferi* s.l. species from the dura of perfused infected animals, further demonstrating the generalized nature of dura colonization in mice for Lyme disease spirochetes. We did not detect spirochetes in the brains of infected animals by culture, qRT-PCR, or fluorescence microscopy, supporting the view that brain colonization in these animals is rare (Garcia-Monco & Benach, 2013).

Bb_297 burdens in the dura mater peaked at day 7 post-infection, and by days 14-28 were quickly reduced to low levels similar to that reported during late disseminated infection (Divan et al., 2018). Intriguingly, this early peak and clearance of spirochetes more closely resembles that reported in blood, rather than other peripheral tissues where burdens peak between 2-3 weeks post-infection and remain relatively higher throughout infection (Aranjuez, Kuhn, Adams, & Jewett, 2019; S. Barthold et al., 1991; Halpern, Jain, & Jewett, 2013; Ma et al., 1998). This rapid reduction of spirochete numbers in the dura mater suggests immune pressures similar to those seen in the blood, and was at least in part due to an adaptive immune response as burdens were significantly higher in SCID mice beyond day 7 post-infection. It has been proposed that the ability of *B. burgdorferi* to persist in peripheral tissues in the face of a strong anti-*Borrelia* immune response is due to tissue-specific protective niches established by host ECM molecules including decorin with high affinity binding to the *B. burgdorferi* surface (Liang, Brown, Wang, Iozzo, & Fikrig, 2004). Given the high levels of decorin, fibronectin, and multiple collagen types found in the dura mater, it is perhaps surprising that this tissue does not serve as such a strong protective niche. Nonetheless, the ability of *B. burgdorferi* to persist at low levels in this tissue for at least 75 days post-infection compared to undetectable levels in the blood during persistent infection may be due to bacterial interactions with these host proteins.

### *Borrelia* colonization of the dura mater is associated with a localized inflammatory response

The dura mater possesses fenestrated blood vessels, lymphatic drainage, and a high density of resident immune cells including dendritic cells (DC), mast cells (MC), innate lymphocytes (ILCs), meningeal macrophages, T cells, and B cells capable of supporting a robust immune response (Rua & McGavern, 2018). Indeed, colonization of the dura mater by Bb_297 was associated with an influx of primarily monocytic leukocytes including T cells. Moreover, widespread changes in gene expression including an inflammatory cytokine and chemokine profile consistent with *B. burgdorferi*-stimulated TLR activation and NF-κB signaling was observed in the dura mater that was sustained up to 8 weeks post-infection. In addition to this inflammatory cytokine response, a transient interferon response was demonstrated that was not dependent on T cells or B cells, consistent with previous studies in *B. burgdorferi*-infected mice (Crandall et al., 2006; Miller et al., 2008). An early interferon response has been shown to be associated with spirochete dissemination, and contributes to murine arthritis pathology as well as affecting the cellular composition of lymph nodes in infected mice (Crandall et al., 2006; Hastey, Ochoa, Olsen, Barthold, & Baumgarth, 2014; Miller et al., 2008; Petzke et al., 2016). It is tempting to speculate that this robust inflammatory response in the dura mater could prime the animal for an inflammatory environment in the leptomeninges and the brain. Although we did not examine the CSF response in this study, a number of genes upregulated in the dura mater of infected mice have been reported to be elevated in the CSF of Lyme neuroborreliosis patients including cytokines/chemokines as well as the matrix metalloproteinases MMP3 and MMP9 (Kirchner et al., 2000; Pietikainen et al., 2016; Yushchenko et al., 2000). Of these, the B cell chemokine Cxcl13 has become of particular interest due to its strong association with Lyme neuroborreliosis compared to healthy controls and patients with other neuroinflammatory diseases (Pietikainen et al., 2016; Wagner, Weis, Kubasta, Panholzer, & von Oertzen, 2018). Co-culture experiments have demonstrated that *B. burgdorferi* induces Cxcl13 production in human dendritic cells and murine synovial cells(X. Wang et al., 2008) (Narayan et al., 2005). A recent study reported that type I interferon induced Cxcl13 production in platelet-derived growth factor receptor α (PDGFRα)+ pulmonary fibroblasts, but not hematopoietic cells, epithelial cells, or endothelial cells in response to Influenza A virus leading to recruitment of B cells (Denton et al., 2019). The presence of dendritic cells and fibroblasts combined with the induction of a strong interferon response in the dura mater implicates either of these cell types as potential sources of the increase in dura *cxcl13* transcript in mice infected with *B. burgdorferi*. Despite the strong induction of *cxcl13* in the dura mater, we did not observe the presence of ectopic germinal centers or an increase in mRNA expression of the genes for the B cell marker *cd19* or the Cxcl13 receptor *cxcr5* at this early timepoint; however, this may be due to temporal or technical limitations of our experimental design.

An interesting finding of this study was the increase in leukocyte influx associated with vascular hemorrhage in the dura mater, cerebral meninges, ventricles, and choroid plexus of mice infected with Bb_297. Although this hemorrhaging may have happened peri- or post-mortem, the fact that this phenomenon was not seen in uninfected animals suggests a decrease in vascular integrity in response to infection. We observed decreased expression of several ECM components important for vascular integrity in the dura mater at day 7 post-infection including BM-associated collagens, laminins, and other structural proteins in addition to increased expression of matrix metalloprotease genes *mmp3* and *mmp9* (Murakami & Simons, 2009; X. Wang & Khalil, 2018). Additionally, we found the endothelial adhesion molecule VE-cadherin gene *cdh5* that is normally expressed in blood and lymphatic vessels of the dura mater was significantly decreased by 44% (p = 3.7e-07), although this did not reach our pre-determined cut-off for differential expression status (Castro Dias, Mapunda, Vladymyrov, & Engelhardt, 2019). Nonetheless, this finding in conjunction with altered expression of ECM components and MMPs provide a potential mechanism for decreased vascular integrity seen in the dura mater of infected mice (Murakami & Simons, 2009; X. Wang & Khalil, 2018). The *B. burgdorferi*-induced breakdown in vascular integrity in the dura mater may explain the rapid accumulation of spirochetes and immune cells during early infection. Further effects on vascular integrity in other tissues or barrier functions in the meninges that could contribute to CNS pathology are yet to be determined; however, vascular leakage in the dura mater alone could lead to symptoms sometimes associated with Lyme disease in humans, including headache (Levy, Labastida-Ramirez, & MaassenVanDenBrink, 2019).

### Mice infected with *B. burgdorferi* exhibit a sterile immune response in the brain parenchyma during early infection

Perhaps our most intriguing finding was evidence of an immune response in the brain cortex and hippocampus, despite the absence of spirochete detection in the parenchyma by molecular, microscopic, or bacterial culture methods. To our knowledge, this is the first description of a tissue-specific sterile immune response to *B. burgdorferi* infection in mice. Gene expression changes in the brain were predominantly upregulation of ISGs. The role for an interferon response in the brain is not known; however, it is noteworthy that both type I and type II interferon can contribute to inflammation in murine models of Lyme arthritis and carditis (Miller et al., 2008; Olson et al., 2009). Specifically, we demonstrated upregulation of genes involved in antigen presentation by both MHCI and MHCII in both cortex and hippocampus of infected mice, as well as increased expression of the gene coding for CD11c in the cortex. No difference was seen for the marker of astrogliosis gene *gfap*. These differences may indicate changes in the immune activation phenotype of resident microglia or an influx of peripheral immune cells, although our histopathologic analysis did not detect the presence of infiltrating leukocytes in the parenchyma. Notably, MHC II expression has been linked to many neurodegenerative diseases in humans and mouse models, and increased CD11c has been demonstrated in transcriptomic studies from both total cortex and acutely isolated microglia in several mouse models of neurodegenerative disease (Schetters, Gomez-Nicola, Garcia-Vallejo, & Van Kooyk, 2017). Interestingly, increased CD11c expression by microglia has been linked to decreased expression of pro-inflammatory cytokines (Wlodarczyk, Lobner, Cedile, & Owens, 2014). Although the pathologic consequences of the observed changes in the brain parenchyma are not clear, these data demonstrate that the brain is not simply a naïve bystander during *B. burgdorferi* infection of laboratory mice.

### Conclusion

Overall, the findings reported in this study are significant, as the lack of a tractable animal model has hindered our understanding of host-pathogen interactions in the CNS during *B. burgdorferi* infection. Our results describe a model system that will allow for future studies evaluating the bacterial, host, and environmental factors that can contribute to the severity of CNS involvement during *B. burgdorferi* infection, as well as evaluating potential novel prophylactic and therapeutic interventions for this important disease.

## MATERIALS AND METHODS

### Bacterial strains and culture conditions

Low passage *B. burgdorferi* isolate 297 strain CE162 (Bb_297) and GFP expressing isogenic mutant Bb914 (Bb_297-GFP) were obtained as gifts from Melissa Caimano and Justin Radolf (Caimano, Eggers, Hazlett, & Radolf, 2004; Dunham-Ems et al., 2009). *B. burgdorferi* strain B31 clone MI-16 was obtained as a gift from Brian Stevenson (Miller, von Lackum, Babb, McAlister, & Stevenson, 2003). *B. mayonii* strain MN14-1539 was obtained as a gift from Jeanine Peterson (Pritt et al., 2016). *B. garinii* strain Ip90 was obtained as a gift from Troy Bankhead (Kriuchechnikov et al., 1988). To confirm the presence of plasmids that were required for infectivity, plasmid content for each strain of *B. burgdorferi* was analyzed by multiplex PCR with primers specific for regions unique to each plasmid, as previously described (Bunikis, Kutschan-Bunikis, Bonde, & Bergstrom, 2011; Xiang et al., 2017). Prior to all animal infections, spirochetes were cultured to mid-log phase in BSK-II medium at 37°C, 5% CO2, and quantified by dark field microscopy using a Petroff-Hausser chamber (Barbour, 1984).

### Animal infections

All animal infections were carried out in accordance with approved protocols from the Institutional Animal Care and Use Committee (IACUC) of the University of North Dakota (Animal Welfare assurance number A3917-01). For all experiments, 6-8 week‐old female mice were used, and housed in groups unless otherwise noted.

For infections by needle inoculation, animals were placed under anesthesia using isoflurane inhalation, followed by intradermal inoculation with 100 uL of BSK-II medium containing the indicated challenge dose and strain of *Borrelia* (infected animals), or medium alone (uninfected controls). Inoculation site included the dorsal thoracic midline, dorsal lumbar midline, or footpad, as indicated.

For infections by tick transmission, larval *Ixodes scapularis* ticks were obtained from BEI Resources (Manassas, VA). Four weeks post-infection by needle inoculation, mice were anesthetized with isoflurane and briefly placed in a tube with naïve larval ticks. To facilitate long-term tick attachment, the mice were placed in individual restrainers overnight. The following day, mice were released and placed on a wire rack suspended over water and provided with food and water ad libitum. Replete ticks were collected daily, stored at 23°C with 12-hour light/dark cycles, and allowed to molt to nymphs. For infection of mice by tick transmission, infected nymphs were applied to naïve mice and allowed to feed as above (10 ticks per mouse). Replete nymphal ticks were collected, crushed, and cultured in BSK medium to confirm infection. The number of recovered infected nymphs ranged from 2-7 per mouse for transmission experiments.

Infections were confirmed in mice by collecting ~80 μL blood from the saphenous vein at day 7 post-infection and cultured in BSK‐II supplemented with 20‐μg ml−1 phosphomycin, 50‐μg ml−1 rifampicin, and 2.5‐ μg ml−1 amphotericin‐B. Ear tissue was also isolated and cultured at time of sacrifice of all animals. Dark‐field microscopy was used to confirm the presence of viable spirochetes for each cultured blood/tissue sample.

### Immunohistochemistry, epifluorescence, and confocal imaging

Samples were stained for imaging of spirochetes, T cells, and endothelial vessels as previously described (Divan et al., 2018; Louveau, Filiano, & Kipnis, 2018). Briefly, each sample was post-fixed in 4% paraformaldehyde (PFA) for 24h at 4°C. Samples were permeabilized in 0.1% Triton X-100, washed 3 times, and serum-blocked in 2.5% goat serum/PBS containing 1:100 dilution of Fc block (CAT # 553142; BD Biosciences, San Jose CA). For *B. burgdorferi* staining, each sample was incubated in 1:100 dilution of rat anti-mouse unconjugated monoclonal anti-CD31 IgG (BD; CAT # 550274), and 1:50 dilution biotinylated rabbit anti-*B. burgdorferi* polyclonal IgG (Invitrogen; CAT# PA1-73007; Thermo-Fisher Scientific, Waltham, MA) at 4°C overnight. On the following day, the samples were washed, and stained with 1:100 dilution of Alexa 555 goat anti-rat polyclonal IgG (Invitrogen; CAT # A-21434), and 1:200 dilution of Alexa 488 streptavidin (Invitrogen; CAT # S11223) for 1 hour at room temperature, covered from light. For CD3 staining, each sample was primary stained using 1:200 dilution of rabbit unconjugated polyclonal anti-CD3 IgG (CAT # ab5690; Abcam, Cambridge, MA), or an equivalent concentration of rabbit unconjugated anti-mouse polyclonal IgG as an isotype control (Abcam; CAT # ab37415). Secondary staining was performed using 1:600 dilution of goat Alexa 488 polyclonal anti-rabbit IgG (Abcam; CAT # ab150081). CD31 staining was performed as described above. For all samples, secondary antibody staining alone was done as a negative control for non-specific signal, and spleen sections were stained as positive control for CD3 binding as previously described (Divan et al., 2018). After antibody staining, samples were incubated in PBS containing 1uM TOPRO-3 nuclear stain for 10 minutes, followed by 2 more washes. Each sample was placed onto a positively charged glass slide and mounted using VECTASHIELD antifade mounting medium (CAT # H-1200; Vector Labs, Burlingame, CA) and gridded coverslip (Electron Microscopy Sciences, Hatfield, PA).

Spirochetes and CD3+ cells stained with Alexa 488 secondary antibody were identified from separate samples by epifluorescence based on morphology and positive signal in the FITC channel using an Olympus BX-50 (Olympus; Center Valley, PA) at 200x magnification as previously described (Divan et al., 2018). Bacteria and CD3+ cells were quantified by manual counting in a blinded fashion. Samples with more than 1000 positive events were counted by systematic random sampling using the gridded coverslips, counting every tenth grid square (McArt et al., 2009). Initial pilot experiments revealed no significant bias in distribution of spirochetes with regards to distance from dural sinuses that would impact the accuracy of systematic counts (not shown). Images were acquired from representative samples above using a Zeiss LSM 510 confocal microscope (Ziess-US; White Plains, NY), and data was collected using Olympus Cell Sens software followed by image processing using Fiji (Divan et al., 2018; Schindelin et al., 2012).

### Intravital microscopy

Skullcaps with attached dura were isolated by craniotomy from mice infected with Bb_297-GFP at day 7 post-infection. Freshly isolated skullcaps were inverted and immobilized to a 100mm X 15mm petri dish, and the exposed dura was covered in PBS to provide a medium for immersion of the objective lens. Imaging was immediately performed in real time using an Olympus FV1000 MPE basic upright multiphoton laser scanning microscope equipped with a tunable MaiTai DS IR laser (690-1040 nm range).

Images were acquired using an Olympus XLPLN 25X, 1.05NA water immersion lens with zoom set to 3.0 and the IR laser tuned to 910 nm. Emission wavelengths of 420-460 nm (violet; second harmonic generation) and 495-540 nm (green: GFP) were used to image connective tissue and the bacteria respectively. Images were acquired using continuous frame capture at 5.43 seconds per frame for 40 frames per image sequence. Image resolution was 165 nm/pixel. Image processing, analysis, and video construction was done using Fiji (Schindelin et al., 2012).

### Histopathology

Skullcaps with attached dura were isolated by craniotomy followed by immediate fixation in 4% PFA for 24h at 4°C. Fixed samples were decalcified in 0.4M EDTA in PBS for 48h at room temperature followed by serial dehydration in 10%/20%/30% sucrose, frozen in Tissue-Tek® OCT (CAT # 4583; Sakura Finetek USA, Torrence, CA), and cut on a cryostat in 6μm coronal sections. Representative sections from each sample (every 35th 6μm section) were stained with haematoxylin and eosin for evaluation by light microscopy. Sections were scored on an increasing scale of 0-3 as follows: No inflammatory cell infiltrate (score = 0); Perivascular infiltrates associated with hemorrhage (score = 1); Perivascular infiltrates without hemorrhage (score = 2); Meningitis (<10% meningitis; score = 3). All histopathology scores were determined by an ACVP board certified veterinary pathologist who was masked to the identity of the samples.

Coronal brain sections (10 μm) containing both cerebral cortex and hippocampus from the same animals were processed in a similar manner without decalcification step, and scored using criteria identical to dura samples.

### RNA isolation

For gene expression analysis, mice were anesthetized using isofluorane and perfused transcardially with 4 mL PBS followed by 4 mL RNAlater™ (Invitrogen; CAT # AM7020) using a peristaltic pump at a flow rate of 0.8mL/min. Dura, heart, and joint tissues were isolated as previously described and immediately snap‐frozen in liquid nitrogen prior to storage at −80°C. Brains were removed and stored in 4 mL RNAlater™ at 4°C for < 1 week prior to processing. A 2mm thick coronal slice of each brain was manually dissected under cold RNAlater™ to isolate cortex and hippocampus from surrounding regions including removing the meninges and corpus collosum. Isolated cortex and hippocampus were snap-frozen in liquid nitrogen and stored at −80°C.

Tissues from 3 separate animals were combined per sample prior to RNA extraction, and represented a single biological replicate. Frozen tissues were ground under liquid nitrogen, added to 1 ml of pre-warmed (65°C) TRIzol reagent (Invitrogen; CAT # 15596026), and frozen at −80°C overnight. Trizol suspensions were thawed at room temperature, and RNA was isolated using the Direct-zol RNA minikit (CAT # R2052; Zymo Research, Irvine CA) according to the manufacturer’s instructions. RNA concentration was determined by a Qubit 2.0 fluorometer (Life Technologies; Carlsbad, CA), and RNA integrity was verified by microfluidic-based capillary electrophoresis with an Agilent 2100 Bioanalyzer (RNA integrity number [RIN] ≥ 8.5 for all samples).

### RNA sequencing

cDNA libraries were prepared from 250ng of purified input RNA using the NEBNext Ultra II kit (CAT#E7770S) with Poly(A) mRNA Magnetic Isolation Module (CAT#E7490S) and index PCR primers (CAT #s E7335, E7500) (New England Biolabs; Ipswich MA) according to the manufacturer’s instructions. Library concentration was assessed with a BioTek Gen5 Wellplate reader with the Quant-iT PicoGreen dsDNA Assay kit (Thermo Fisher; CAT # P11496), and analyzed on the Bioanalyzer to ensure appropriate size distributions and rule out adaptor contamination.

The indexed cDNA libraries were pooled and 150 bp paired-end reads were sequenced on two lanes using the Illumina HiSeq 4000 (Novogene; San Diego, CA). Demultiplexed fastq files from the two sequencing runs were combined for each sample, and read quality confirmed using FASTQC v0.11.2 prior to analysis.

For the analysis of transcriptome sequencing (RNA-seq) data, adapters were removed from the sequencing reads by Trimmomatic v0.32 (Bolger, Lohse, & Usadel, 2014). Reads were aligned to the murine genome (mm10) using STAR v2.7.1a with 2-pass mapping (Dobin et al., 2013). Fragments were assigned to genes using featureCounts v1.6.4 (Liao, Smyth, & Shi, 2014). Differential expression analysis was performed using DESeq2 v1.24.0 (Love, Huber, & Anders, 2014). Genes were considered to be differentially expressed in infected samples compared to control samples at a false discovery rate (padj) of ≤0.05, basemean>20, fold-change >1.5.

Functional analysis (gene ontology) of upregulated or downregulated genes was performed using clusterProfiler v3.12.0 (Yu, Wang, Han, & He, 2012). KEGG pathway enrichment analysis from all DEGs was performed using Signaling Pathway Impact Analysis (SPIA) v2.36.0 with Bonferroni correction and a significance threshold of pGFWER ≤ 0.05 (Tarca et al., 2009).

### Quantitative reverse transcriptase PCR (qRT-PCR)

cDNA was generated from 500ng of purified RNA using Superscript® IV First-Strand Synthesis System with included RnaseH treatment (Invitrogen; Cat # 18091050). Quantitative PCR for the *B. burgdorferi flaB* gene was performed in 20μL reactions with 12.5ng cDNA, gene-specific primers, and internal fluorescent probes using Bio-Rad SsoAdvanced™ Universal probes Supermix (CAT # 1725281; Bio-Rad, Hercules, CA) as previously described (Casselli, Crowley, Highland, Tourand, & Bankhead, 2019). Absolute copy numbers were interpolated for each sample in triplicate using standard curves. Host gene expression was determined with individual PCR primer sets (*gapdh*, CAT#QT01658692; *tnfa*, CAT#QT00104006; *gbp2*, CAT#QT00106050; *cxcl10*, CAT#QT00093436; Qiagen USA, Germantown, MD) using Bio-Rad SsoAdvanced™ Universal SYBR® Green Supermix (Bio-Rad; CAT #1725274). Relative changes in gene expression were compared between infected and control animals using the 2-^ΔΔCt^ method with *gapdh* as a housekeeping control.

### Statistical analysis

Statistical analysis of RNA sequencing data is described above in the “RNA sequencing” subsection of Materials and Methods. Statistical tests used to compare means for all other experiments were performed using Sigmaplot v11.0 (Systat Software; San Jose, CA) and are described in the relevant figure legends. For microscopy counting experiments, sample sizes were calculated a priori using G*Power v 3.1.9.7 using the following input parameters: α = 0.05; Power = 0.8; sample N allocation ration = 1; effect size = 5. For qRT-PCR confirmation of RNA-seq results, n = 4 was used to maintain consistency with RNA-seq experiment.

### Data visualization

Data generated from DNA and RNA sequencing analyses were visualized with R v.3.3.0 (https://www.R-project.org/) using the following packages: clusterProfiler v3.12.0 for GO term analysis and cnet plots (Yu et al., 2012); Pathview for KEGG pathway DEG visualization (Luo & Brouwer, 2013); ggplot2 for volcano plots (Wickham, 2016); NMF v0.21.0 for heatmaps (Gaujoux & Seoighe, 2010). All other plots were generated using Sigmaplot v11.0 (Systat Software).

### Accession number

Sequences have been deposited in the NCBI GEO sequence read archive database under accession number XXXXXX.

## ACKNOWLEDGEMENTS

The authors would like to thank Dr. Bryon Grove and Sarah Abrahamson for technical assistance with intravital and confocal imaging, Beth Ann DeMontigny for assistance with histology sample preparation, and Hannah Huffman and Dr. Bony De Kumar for assistance with RNA sequencing sample preparation and data analysis, respectively. We also thank Dr. Brian Stevenson for critical reading of the manuscript. This work was supported by NIH grants P20GM113123 to Catherine Brissette, and R21AI129522 to Brian Stevenson, Wolfram Zueckert, and Catherine Brissette. Imaging studies and histological services were conducted in the UND Imaging Core and Histology Core facilities, respectively, supported by NIH grant P20GM113123, DaCCoTA CTR NIH grant U54GM128729, and UNDSMHS funds. RNA sequencing library preparation/QC and data analysis were carried out in conjuction with the UND genomics core supported by the National Institute of General Medical Sciences of the National Institutes of Health under Award Number U54GM128729 and Award number 2P20GM104360-06A1.

## LIST OF SUPPLEMENTAL MATERIALS

### Supplemental Movies/Figures

**Movie 1. *B. burgdorferi* in the dura mater are extravascular and motile.**

**Figure S1. Gene expression changes are consistent across biological replicates.**

**Figure S2. Genes upregulated in the dura mater and brain parenchyma represent different gene ontologies.**

**Figure S3. Toll-like receptor signaling is increased in the dura mater in response *to B. burgdorferi* infection.**

**Figure S4. NF-κB signaling is increased in the dura mater in response *to B. burgdorferi* infection.**

**Figure S5. Antigen processing and presentation is increased in the dura mater and brain parenchyma in response *to B. burgdorferi* infection.**

**Figure S6. T cell receptor signaling is increased in the dura mater in response *to B. burgdorferi* infection.**

### Supplemental Tables

**Table S1. Differential expression gene list in the dura mater.**

**Table S2. Differential expression gene list in the cortex.**

**Table S3. Differential expression gene list in the hippocampus.**

