## Supplemental Figures S1-S6 and Movie 1 legend for "A Murine Model of Lyme Disease Demonstrates That *Borrelia burgdorferi* Colonizes the Dura Mater and Induces Inflammation in the Central Nervous System"

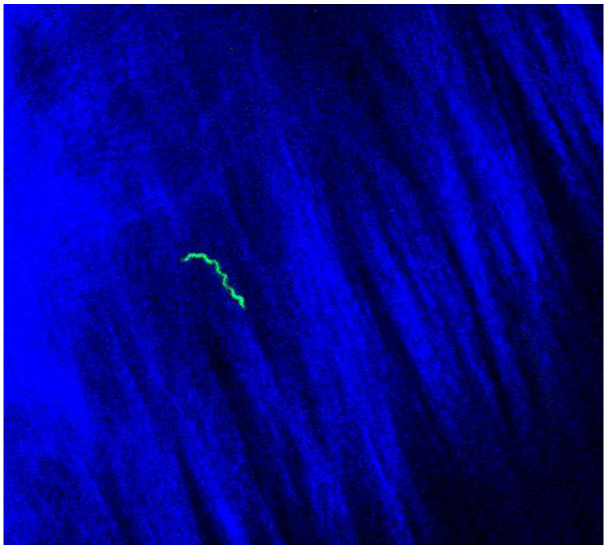

**Movie 1. *B. burgdorferi* in the dura mater are extravascular and motile.** Multiphoton image series of dura mater from C3H mouse after infection with GFP-Bb_297 for 7 days. *B. burgdorferi* is shown in green; second harmonics shown in blue. Two minute movie. Imaging parameters: Wavelength = 910 nm, pixel resolution = 135. Note this is a representative image. See Movie 1 for full movie.

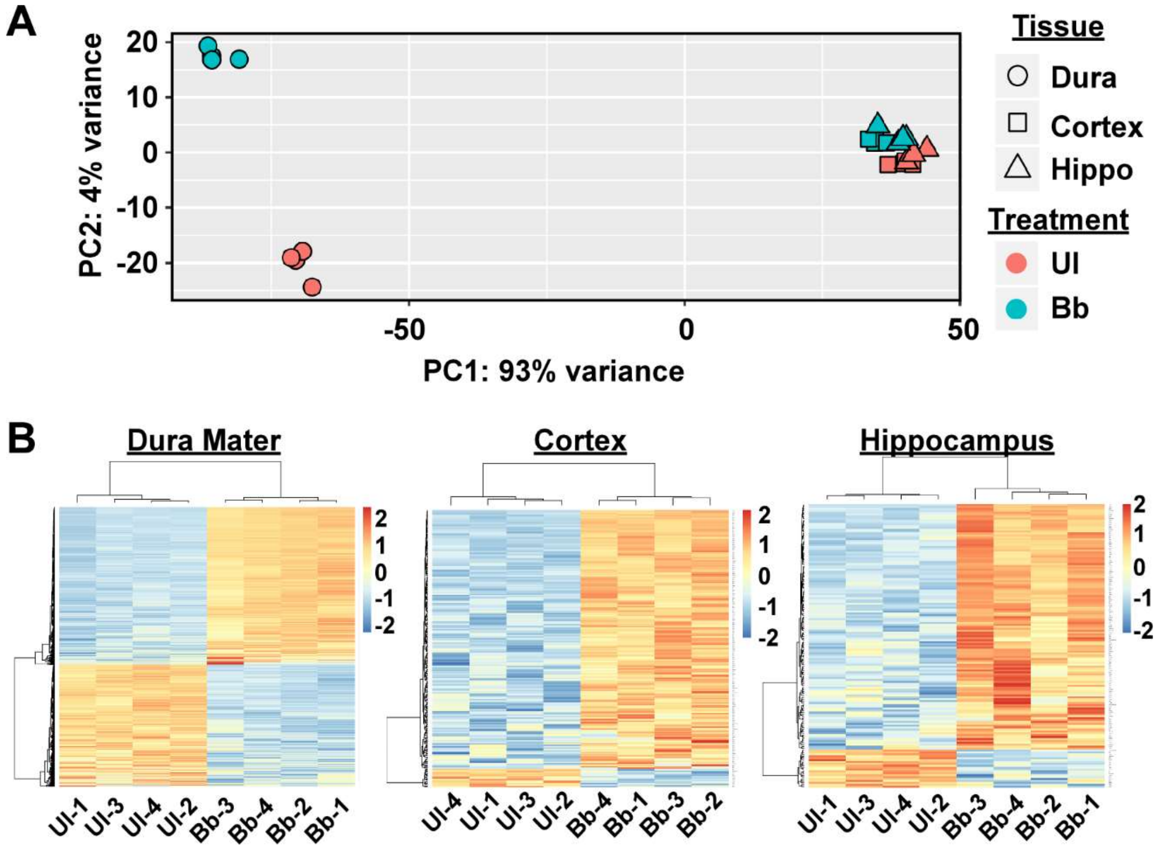


**Figure S1. Gene expression changes are consistent across biological replicates. A.** Principal component analysis of all samples used in this study. Samples are color-coded by treatment (uninfected (UI) vs 7 day Bb_297 infected (Bb)); while tissues are denoted by symbol shape as shown in the legend. **B.** Hierarchical clustering of individual samples from RNA-seq data. Columns represent individual samples, while rows represent individual DEGs. Colors represent euclidean distances from the regularized log-transformed counts (rlog) generated using DESeq2 for all DEGs. Separate heatmaps are displayed for each tissue as indicated in the titles. Treatment groups (uninfected, UI1-4; 7 day infected, Bb1-4) cluster together for each tissue, and show similar profiles of upregulated/downregulated genes in response to infection.

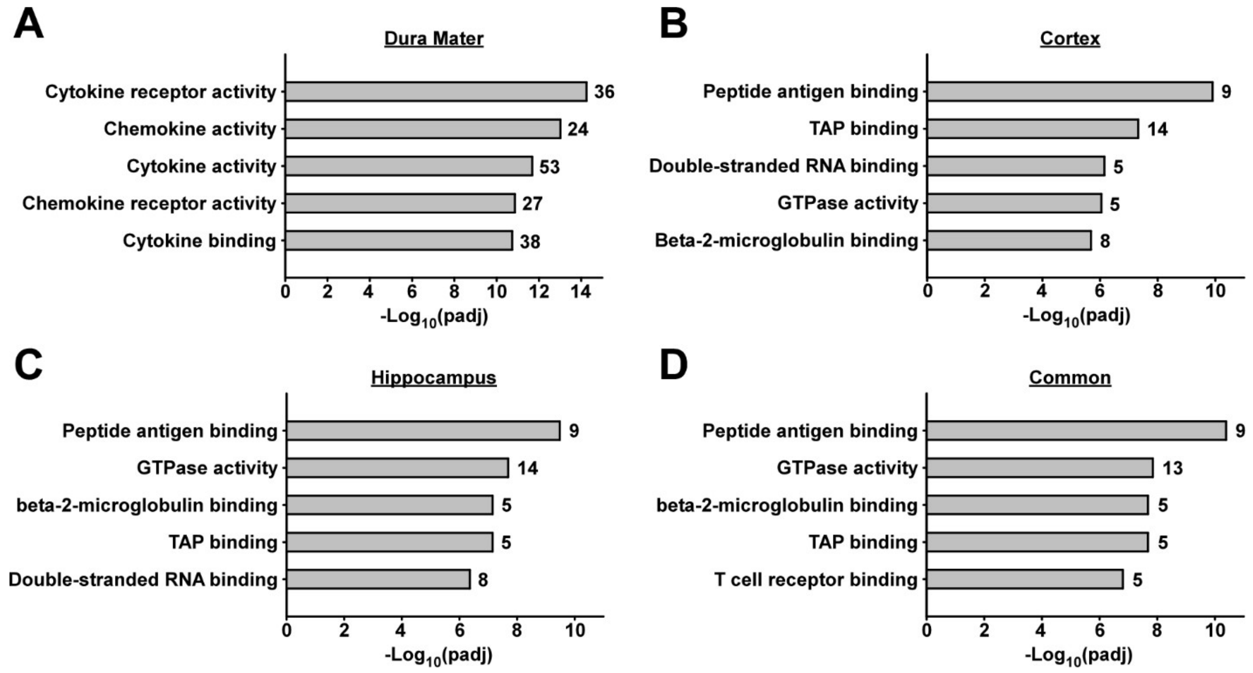


**Figure S2. Genes upregulated in the dura mater, cortex, and hippocampus represent different gene ontologies.** Top five most enriched molecular function gene ontology (GO) terms from upregulated genes in the dura mater **(A)**, cortex **(B)**, hippocampus **(C)**, and genes commonly upregulated in all three tissues **(D)**. Numbers to the right of horizontal bars show the number of upregulated DEGs associated with each term. Bar size represents significance of enrichment (-log(padj)).

## A

| Name | KEGG ID | Pathway Size (# Genes) | Tissue | # DEGs | pGFWER | Status |
|---|---|---|---|---|---|---|
| Toll-like | | | Dura | 38 | 1.30E-06 | Activated |
| receptor | mmu04620 | 101 | Cortex | 4 | 1 | n.s. |
| signaling | | | Hippo | 2 | 1 | n.s. |

## B

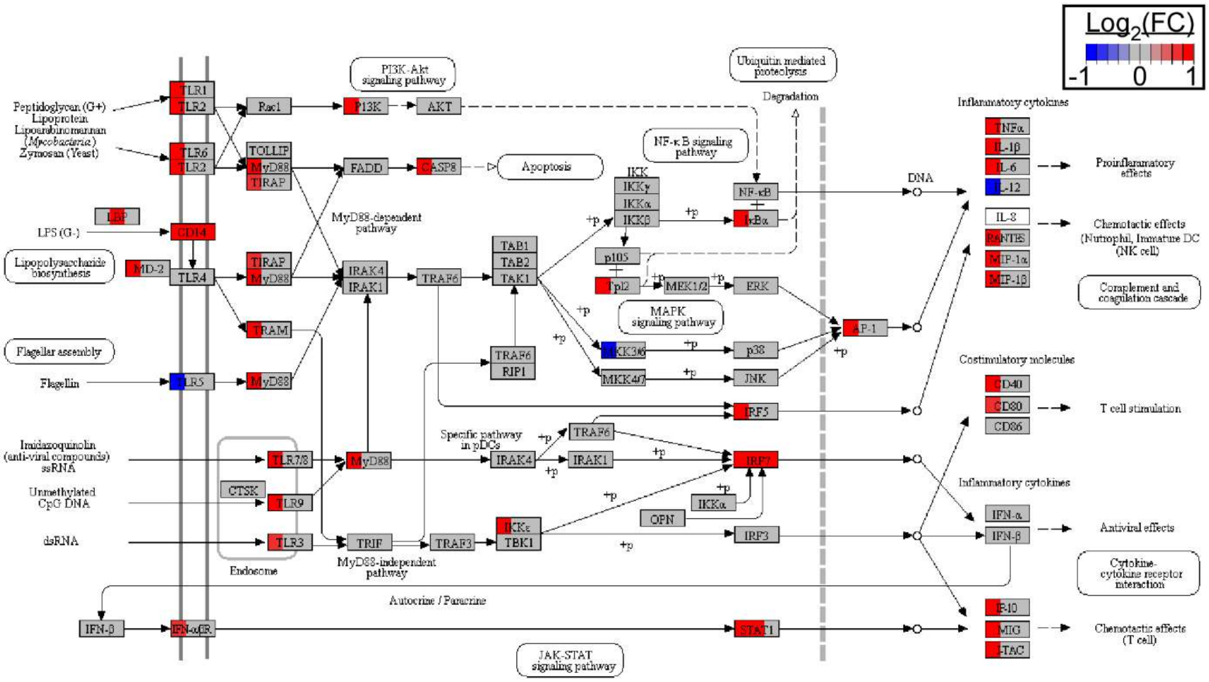


**Figure S3. Toll-like receptor signaling is increased in the dura mater in response *to B. burgdorferi* infection. A.** Summary of Signaling Pathway Impact Analysis (SPIA) for TLR signaling (mmu04620). Table shows number of DEGs for each tissue (dura, cortex, hippocampus) within the pathway, as well as the activation status of the pathway (n.s. = not significant). pGFWER represents the false discovery rate after Bonferroni correction. **B.** DEGs were mapped onto the KEGG pathway as rendered using Pathview (ref). Pathway gene products (such as receptors, adaptors and enzyme proteins) are represented as rectangles, with interaction shown as arrows. Rectangles are color coded by $log_2$(fold-change) from RNA-seq datasets (infect vs. uninfected), with the left one-third of the rectangle representing DEG status in the dura mater, the center representing DEG status in the cortex, and the right one-third representing DEG status in the hippocampus. Color scale is shown in the legend. Upregulation of most TLR signaling genes is restricted to the dura mater following infection.

## A

| Name | KEGG ID | Pathway Size (# Genes) | Tissue | # DEGs | pGFWER | Status |
|---|---|---|---|---|---|---|
| NF-κB signaling | mmu04064 | 96 | Dura | 46 | 1.15E-16 | Activated |
| | | | Cortex | 6 | 0.09 | n.s. |
| | | | Hippo | 4 | 0.37 | n.s. |

## B

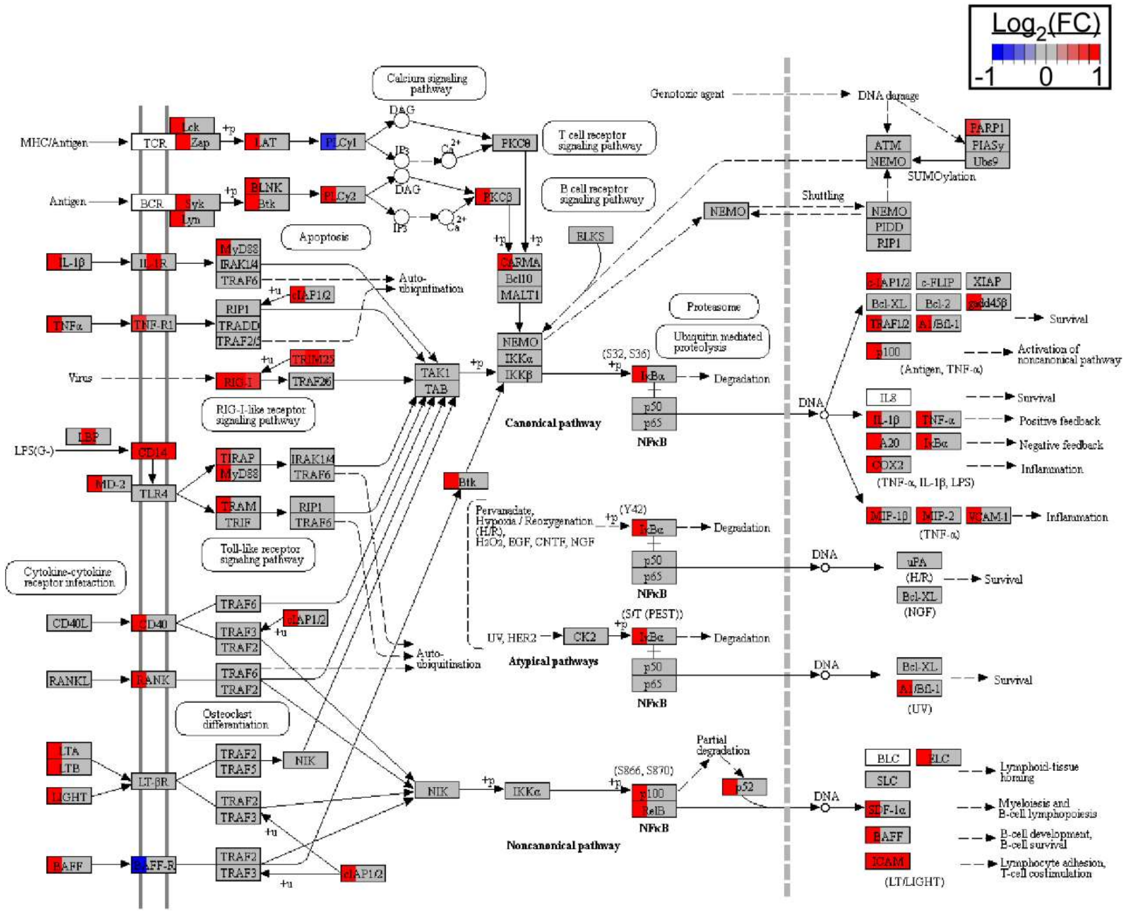


**Figure S4. NF-κB signaling is increased in the dura mater in response *to B. burgdorferi* infection. A.** Summary of Signaling Pathway Impact Analysis (SPIA) for NF-κB signaling (mmu04064). Table shows number of DEGs for each tissue (dura, cortex, hippocampus) within the pathway, as well as the activation status of the pathway (n.s. = not significant). pGFWER represents the false discovery rate after Bonferroni correction. **B.** DEGs were mapped onto the KEGG pathway as rendered using Pathview (ref). Pathway gene products (such as receptors, adaptors and enzyme proteins) are represented as rectangles, with interaction shown as arrows. Rectangles are color coded by $\log_2$(fold-change) from RNA-seq datasets (infect vs. uninfected), with the left one-third of the rectangle representing DEG status in the dura mater, the center representing DEG status in the cortex, and the right one-third representing DEG status in the hippocampus. Color scale is shown in the legend. Upregulation of most NF-κB signaling genes is restricted to the dura mater following infection.

**A**

| Name | KEGG ID | Pathway Size (# Genes) | Tissue | # DEGs | pGFWER | Status |
|---|---|---|---|---|---|---|
| Antigen processing and presentation | mmu04612 | 78 | Dura | 36 | 3.62E-13 | Activated |
| | | | Cortex | 17 | 8.13E-15 | Activated |
| | | | Hippo | 11 | 1.84E-09 | Activated |

**B**

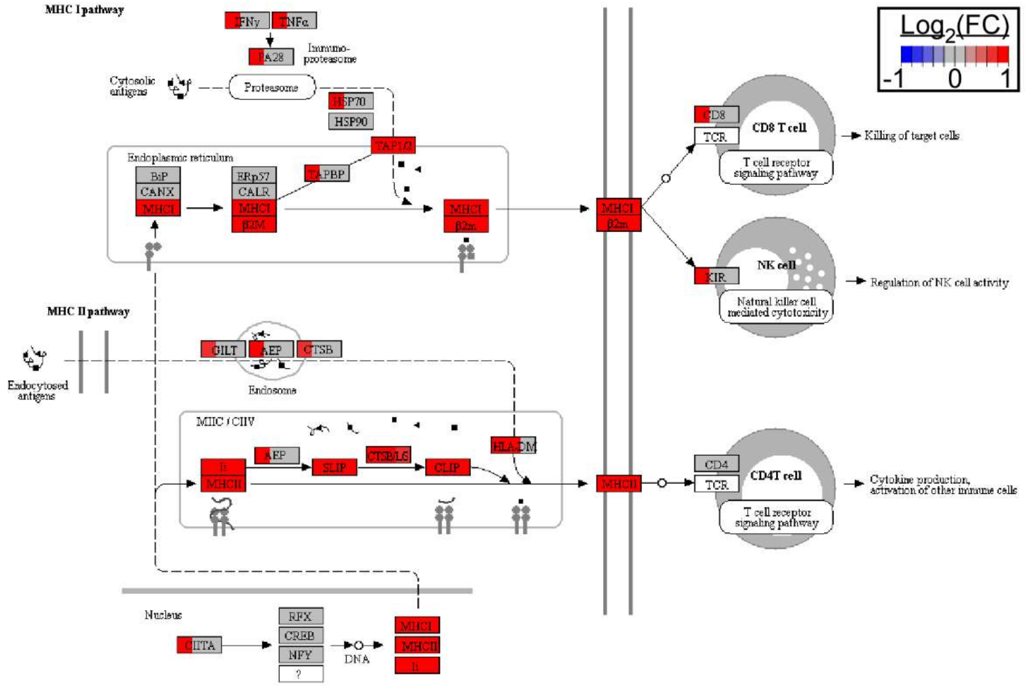


**Figure S5. Antigen processing and presentation is increased in the dura mater and brain parenchyma in response *to B. burgdorferi* infection. A.** Summary of Signaling Pathway Impact Analysis (SPIA) for antigen processing and presentation (mmu04612). Table shows number of DEGs for each tissue (dura, cortex, hippocampus) within the pathway, as well as the activation status of the pathway (n.s. = not significant). pGFWER represents the false discovery rate after Bonferroni correction. **B.** DEGs were mapped onto the KEGG pathway as rendered using Pathview (ref). Pathway gene products (such as receptors, adaptors and enzyme proteins) are represented as rectangles, with interaction shown as arrows. Rectangles are color coded by $log_2$(fold-change) from RNA-seq datasets (infect vs. uninfected), with the left one-third of the rectangle representing DEG status in the dura mater, the center representing DEG status in the cortex, and the right one-third representing DEG status in the hippocampus. Color scale is shown in the legend. Upregulation of most antigen processing/presentation genes is seen in all three tissues following infection.

## A

| Name | KEGG ID | Pathway Size (# Genes) | Tissue | # DEGs | pGFWER | Status |
|---|---|---|---|---|---|---|
| T cell receptor signaling | mmu04660 | 109 | Dura | 37 | 8.19E-05 | Activated |
| | | | Cortex | 1 | 0.97 | n.s. |
| | | | Hippo | 0 | N/A | n.s. |

## B

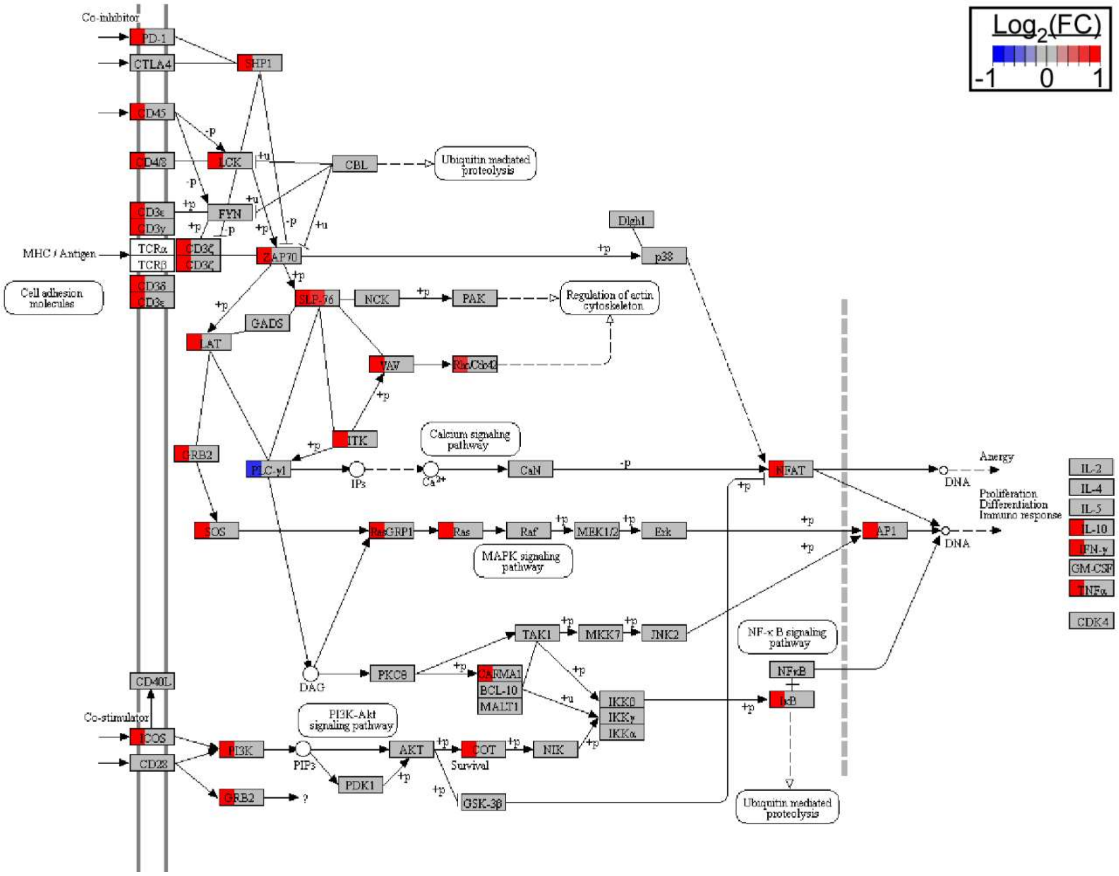


**Figure S6. T cell receptor signaling is increased in the dura mater in response *to B. burgdorferi* infection. A.** Summary of Signaling Pathway Impact Analysis (SPIA) for T cell receptor signaling (mmu04660). Table shows number of DEGs for each tissue (dura, cortex, hippocampus) within the pathway, as well as the activation status of the pathway (n.s. = not significant). pGFWER represents the false discovery rate after Bonferroni correction. **B.** DEGs were mapped onto the KEGG pathway as rendered using Pathview (ref). Pathway gene products (such as receptors, adaptors and enzyme proteins) are represented as rectangles, with interaction shown as arrows. Rectangles are color coded by $\log_2$(fold-change) from RNA-seq datasets (infect vs. uninfected), with the left one-third of the rectangle representing DEG status in the dura mater, the center representing DEG status in the cortex, and the right one-third representing DEG status in the hippocampus. Color scale is shown in the legend. Upregulation of most T cell receptor signaling genes is restricted to the dura mater following infection.
